# A nematode-trapping fungus orchestrates polarity cues, septins, and NOX signaling for trap formation

**DOI:** 10.64898/2026.03.09.710510

**Authors:** Chih-Yen Kuo, Hung-Che Lin, Yun-Zhen Chou, Sheng-An Chen, Hillel Schwartz, Yen-Ping Hsueh

**Author notes:** **Correspondence** Yen-Ping Hsueh Department of Complex Biological Interactions, Max Planck Institute for Biology, Tübingen, Germany.

## Abstract

The ability to reprogram cell morphogenesis in response to environmental cues is fundamental to microbial adaptation and survival. The nematode-trapping fungus *Arthrobotrys oligospora* exemplifies this plasticity by developing adhesive traps to capture nematodes. This morphological transition involves spatial reorientation of cell polarity, cytoskeletal remodeling, and cell fusion. However, the molecular mechanisms that coordinate these processes remain unclear. Using live-cell imaging, genetics, and functional assays, we demonstrated that cell polarity proteins localize to hyphal tips to direct growth, while actin and septins assemble at the curving inner rim to shape trap architecture. Conserved NADPH oxidases are induced by the presence of nematodes and are required for the recruitment of cell polarity proteins and cytoskeletal factors to promote trap cell fusion. Together, these findings reveal that polarity cues, cytoskeletal organization, and reactive oxygen species signaling are integrated to orchestrate nematode-induced trap development, establishing nematode-trapping fungi as a versatile model to study fungal cell biology.

## Introduction

Fungal survival and pathogenicity depend on the ability to remodel cellular architecture in response to environmental cues (Riquelme et al, 2018). This morphogenetic plasticity enables fungi to switch between growth forms or develop specialized infection structures, such as host-penetrating appressoria (Boyce & Andrianopoulos, 2015; Ryder & Talbot, 2015). Cell polarity is the primary organizer of these transitions (Asnacios & Hamant, 2012; Chiou et al, 2017). In the fission yeast *Schizosaccharomyces pombe,* the cell-end markers Tea1 and Tea4 are delivered by microtubules and anchored by Mod5 to define sites of polarized growth (Martin et al, 2005; Mata & Nurse, 1997; Snaith & Sawin, 2003). In filamentous fungi, this conserved polarity machinery supports continuous apical hyphal extension (Kriegler et al, 2025a; Takeshita et al, 2013). During hyphal extension, chitin synthases (CHSs) are delivered along actin cables to the Spitzenkörper (SPK), where polarized secretion directs localized chitin deposition at the hyphal (Riquelme & Sánchez-León, 2014). While linear growth relies on steady apical extension, complex morphogenesis requires dynamic reorganization of polarity cues and membrane architecture.

The cytoskeleton acts as both a transport track and a scaffold for shape remodeling (Riquelme et al, 2018). Actin filaments form apical patches and contractile rings to mediate endocytosis and septum formation (Berepiki et al, 2011; Berepiki et al, 2010). Septins are a conserved family of GTP-binding proteins that assemble into **filaments and ring-like structures** and are often regarded as a fourth cytoskeletal element in animals and fungi (Mostowy & Cossart, 2012). They preferentially assemble at regions of micron-scale membrane curvature, such as yeast mother–bud necks and hyphal branches (Bridges et al, 2016; Khan et al, 2015). In the rice blast fungus *Magnaporthe oryzae*, septins assemble at the appressorium pore to recruit a toroidal F-actin network, which is anchored to the plasma membrane via phosphoinositide interactions and stabilized by Tea1 (Dagdas et al, 2012). These examples highlight the tight coordination between cell polarity proteins and cytoskeletal dynamics in fungal morphogenesis.

Cell fusion is a fundamental process in fungal growth and development, enabling the establishment of interconnected hyphal networks (Fischer & Glass, 2019). Best characterized during the formation of conidial anastomosis tubes (CATs), this process involves MAP kinase signaling that activates NADPH oxidases (NOX enzymes) (Herzog et al, 2015). These enzymes generate reactive oxygen species (ROS) that act as localized signals to trigger cell wall remodeling and membrane merger at the fusion interface (Read et al, 2012; Roca et al, 2012). NOX-derived ROS serve as essential regulators of fungal development, controlling processes from hyphal branching to sexual reproduction (Tudzynski et al, 2012).

Nematode-trapping fungi (NTF) represent a powerful model for studying fungal morphogenetic plasticity. Under nutrient-limiting conditions, NTF sense nematode-derived signals and shift from a saprophytic to a predatory lifestyle by developing trapping structures to capture nematodes as nutrient source (Nordbring-Hertz et al, 2001). In response to prey, the NTF *Arthrobotrys oligospora* produces adhesive traps through a stereotypic program where specialized hyphae grow, curve, and fuse to form closed loops that generate intricate three-dimensional networks. Signaling pathways governing trap induction have been characterized (Chen et al, 2022; Chen et al, 2021; Hu et al, 2024; Kuo et al, 2024). However, the cellular mechanisms coordinating loop curvature and cell fusion during trap morphogenesis remain largely unresolved, despite evidence that cell-end markers, septin Cdc11, and NoxA play a role in trap formation in *A. oligospora* or *Arthrobotrys flagrans* (Kriegler et al, 2025b; Li et al, 2017; Zhu et al, 2025).

In this study, we asked how *A. oligospora* reprograms its cellular machinery to facilitate the transition to a predatory lifestyle. Using live-cell imaging and functional genetics, we demonstrate that cell polarity proteins and cytoskeletal systems are dynamically reorganized during the transition from linear growth to specialized trap and invasive structures. Our findings reveal that: (1) cell-end markers and CHSs relocalize to direct trap growth; (2) asymmetric actin and septin assembly characterizes the inner rim of developing loops; and (3) NOX enzyme-mediated ROS production serves as a localized cell fusion checkpoint. Together, our study reveals how cell polarity proteins and cytoskeletal regulators coordinate morphogenetic transitions in NTF, providing new insights into the fundamental mechanisms of polarized growth and network assembly in filamentous fungi.

## Results

### Identification and characterization of chitin synthases and cell-end markers in *A. oligospora*

To investigate the molecular basis of polarized growth and trap morphogenesis, we identified gene families involved in chitin synthesis and cell polarity in *A. oligospora*. Seven chitin synthase (CHS) genes were identified in the genome of strain TWF154 (Table EV1) (Yang et al, 2020), representing the seven conserved classes found in filamentous fungi (Fig. 1A). Chs1 to Chs6 share approximately 60% identity with their respective orthologs and exhibit significant internal divergence (Fig. EV1). For example, Chs1 to Chs3 exhibit roughly 45% pairwise identity, while Chs4 to Chs6 share approximately 35% identity. Chs7 is the most divergent member, displaying only 15% to 20% identity to other CHSs (Fig. EV1).

**Figure 1.**
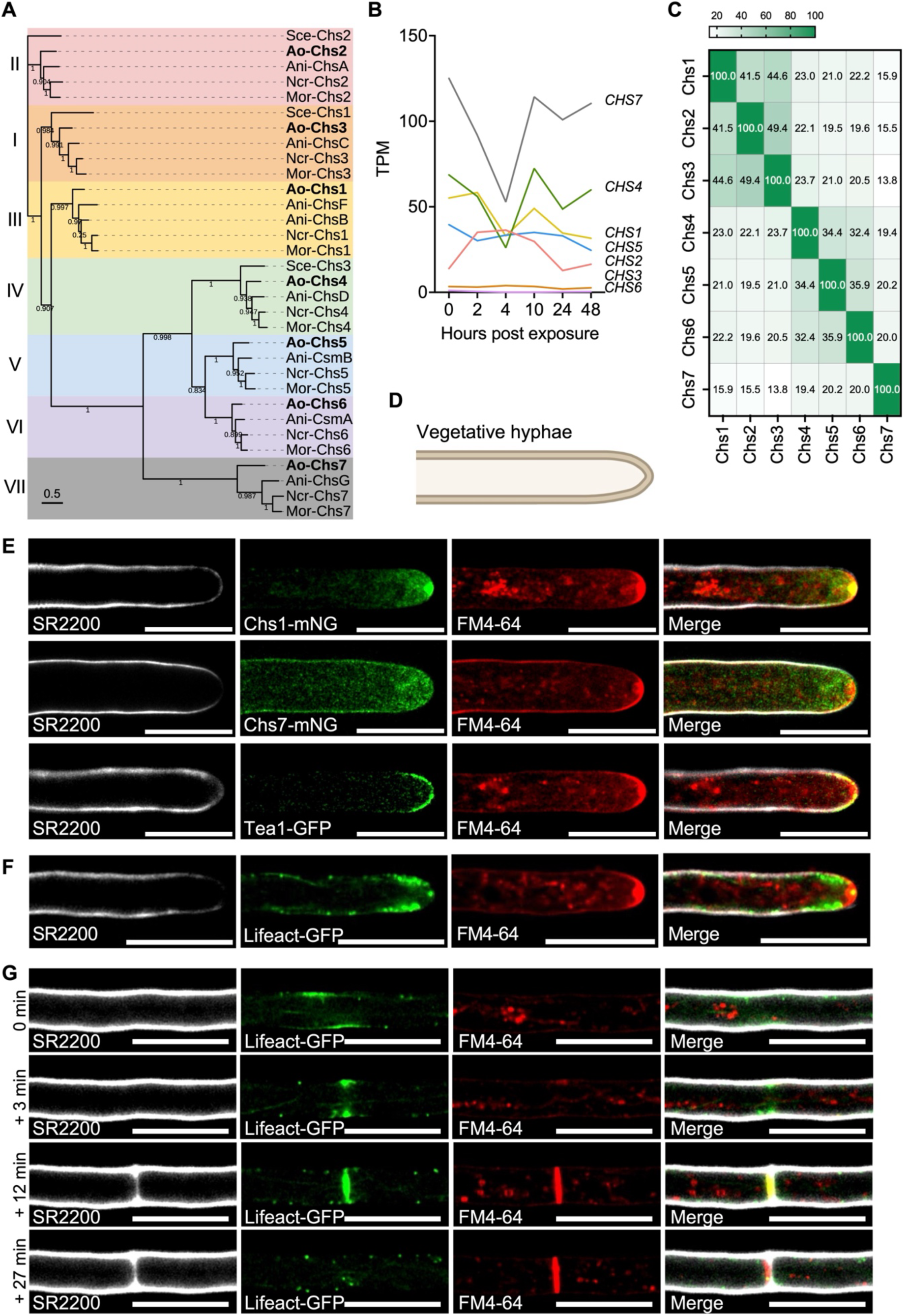
Genomic, transcriptomic, and localization analyses of polarity-related genes in *A. oligospora*. **(A)** Phylogenetic analysis of the fungal chitin synthases Chs1, Chs4, and Chs7. The maximum likelihood tree was constructed using FastTree with protein sequences from *A. oligospora* and orthologs from representative model fungi. Roman numerals on the left denote the CHS classes. Node numbers indicate local support values ranging from 0 to 1. The scale bar represents 0.5 amino acid substitutions per site. Species abbreviations are as follows: Ani, *Aspergillus nidulans*; Ao, *Arthrobotrys oligospora*; Mor, *Magnaporthe oryzae*; Ncr, *Neurospora crassa*; Sce, *Saccharomyces cerevisiae*. **(B)** Transcripts per kilobase million (TPM) values of the differentially expressed *CHS*s in a published RNA-seq time course following exposure of *A. oligospora* to *C. elegans* (Lin et al, 2023). Values represent the average of three independent biological replicates. **(C)** Heatmap displaying pairwise amino acid sequence identity among *A. oligospora* CHSs. The matrix relies on a multiple sequence alignment generated using the T-Coffee algorithm. The color gradient indicates the degree of conservation, where darker shading corresponds to higher identity percentages. Numerical values within the cells represent percent identity for each protein pair. Species abbreviations are consistent with (A). **(D)** Schematic representation of a vegetative hyphal apex. **(E)** The localization of Chs1–mNG, Chs7-mNG, and Tea1–GFP in growing vegetative hyphae. CHSs and Tea1 (green) were enriched at the hyphal tips. **(F-G)** Localization of Lifeact-GFP during vegetative hyphae growth **(F)** and septum formation **(G)**. In all panels, hyphae were co-stained with SR2200 to label cell walls and FM4-64 to visualize the plasma membrane and the Spitzenkörper (SPK). The SPK is visible as a distinct fluorescent spot at the hyphal tip, representing the accumulation of membrane-bound vesicles. Merged images display the SR2200, GFP or mNG, and FM4-64 channels. All scale bars are 10 μm. Images represent three independent biological replicates.

To identify which CHSs are relevant to trap morphogenesis, we examined published *A. oligospora* time-course RNA-seq data following *Caenorhabditis elegans* exposure (Lin et al, 2023). Transcriptional profiling identified *CHS1*, *CHS4*, and *CHS7* as nematode-responsive genes, with transcript levels decreasing at 4 h post-exposure (hpe) and recovering by 10 hpe during trap formation (Fig. 1B). Despite their shared expression profile, these three proteins are highly divergent from each other, sharing only about 20% pairwise identity (Fig. 1C). To examine subcellular localization, Chs1 and Chs7 were C-terminally tagged with mNeonGreen (mNG) under their endogenous promoters. During hyphal extension, Chs1-mNG and Chs7-mNG both localized in the cytoplasm with strong enrichment at the hyphal apex (Fig. 1D and E). Chs1-mNG intensely concentrated as a sharp, central punctum that overlapped with the FM4-64-labeled SPK. In contrast, Chs7-mNG exhibited a diffuse apical distribution. Although present within the apical dome, no significant signal was detected within the central SPK region (Fig. 1E). This spatial differentiation suggests that Chs1 is specifically associated with the central secretory apparatus, while Chs7 is localized to the peripheral apical cytoplasm.

To assess whether conserved fungal cell polarity machinery contributes to trap morphogenesis, we searched the *A. oligospora* genome for homologs of the *S. pombe* cell-end markers. We could identify orthologs of Tea1, Tea2, Tea4, and the membrane anchor Mod5, which we renamed **Tea5** in *A. oligospora,* but found no evidence of the presence of a Tea3 homolog (Table EV1). Phylogenetic analysis demonstrated that these proteins are most closely related to orthologs in *A. nidulans* and the closely related NTF *A. flagrans* (Fig. EV2A-D). Domain architecture analysis revealed high conservation of functional motifs, including Kelch repeats in Tea1, a kinesin motor domain in Tea2, an SH3 domain in Tea4, and a C-terminal CAAX-like prenylation motif in Tea5 (Fig. EV2E).

RNA-seq analysis showed that *TEA1*, *TEA2*, *TEA4*, and *TEA5* are nematode-responsive genes, with peak expression at 10 hpe (Fig. EV2F). To examine Tea1 localization during hyphal extension, we generated a strain expressing Tea1-GFP fusion under its native promoter. Tea1-GFP exhibited a highly polarized distribution and localized predominantly to the apical plasma membrane. Unlike the central punctum observed for Chs1, Tea1-GFP remained confined to the peripheral cortex of the hyphal apex, forming a crescent-like cap that delineates the site of active expansion (Fig. 1E).

### Cytoskeletal networks display distinct localization patterns at the tip and curvature of vegetative hyphae

Actin and septins are cytoskeleton proteins that play essential roles in fungal cell polarity and morphogenesis (Riquelme et al, 2018). To visualize F-actin organization during hyphal extension, Lifeact-GFP was expressed under the constitutive promoter in *A. oligospora*. Live imaging revealed that Lifeact–GFP localized to the SPK and appeared as peripheral puncta at hyphal tips (Fig. 1F; Movie EV1). These puncta represent actin patches that recruit endocytic machinery, suggesting spatial coupling between apical exocytosis and compensatory endocytosis (Galletta & Cooper, 2009). During septation, time-lapse imaging revealed de novo assembly of a medial actin ring at future division sites (Fig. 1G; Movie EV1). The actin ring formed before visible membrane ingression and then constricted as the plasma membrane progressed, as indicated by FM4-64 staining. As septum formation progressed, the actin signal gradually diminished. These dynamics indicate that actin ring assembly precedes and likely drives septal membrane ingression and cell wall deposition during cytokinesis.

Given the role of septins in membrane remodeling and cell morphology (Benoit et al, 2023), we analyzed the *A. oligospora* genome and identified six septin genes (Fig. 2A; Table EV1). Sep3, Sep4, Sep5, and Sep6 belong to the four core septin groups (1a, 2a, 3, 4), while Sep7 and Sep8 belong to the fifth group (5), which is unique to filamentous fungi and protists (Shuman & Momany, 2022). Sequence analyses showed that the core septins are highly conserved, sharing 85 to 90% sequence identity with orthologs in *A. nidulans* and *M. oryzae; whereas Sep7 and Sep8 are much less conserved* (Fig. 2B). Analysis of a previously published time-course RNA-seq dataset (Lin et al, 2023) revealed that the core septins are all up-regulated in response to *C. elegans*, suggesting a role in trap morphogenesis and development (Fig. EV2G).

**Figure 2.**
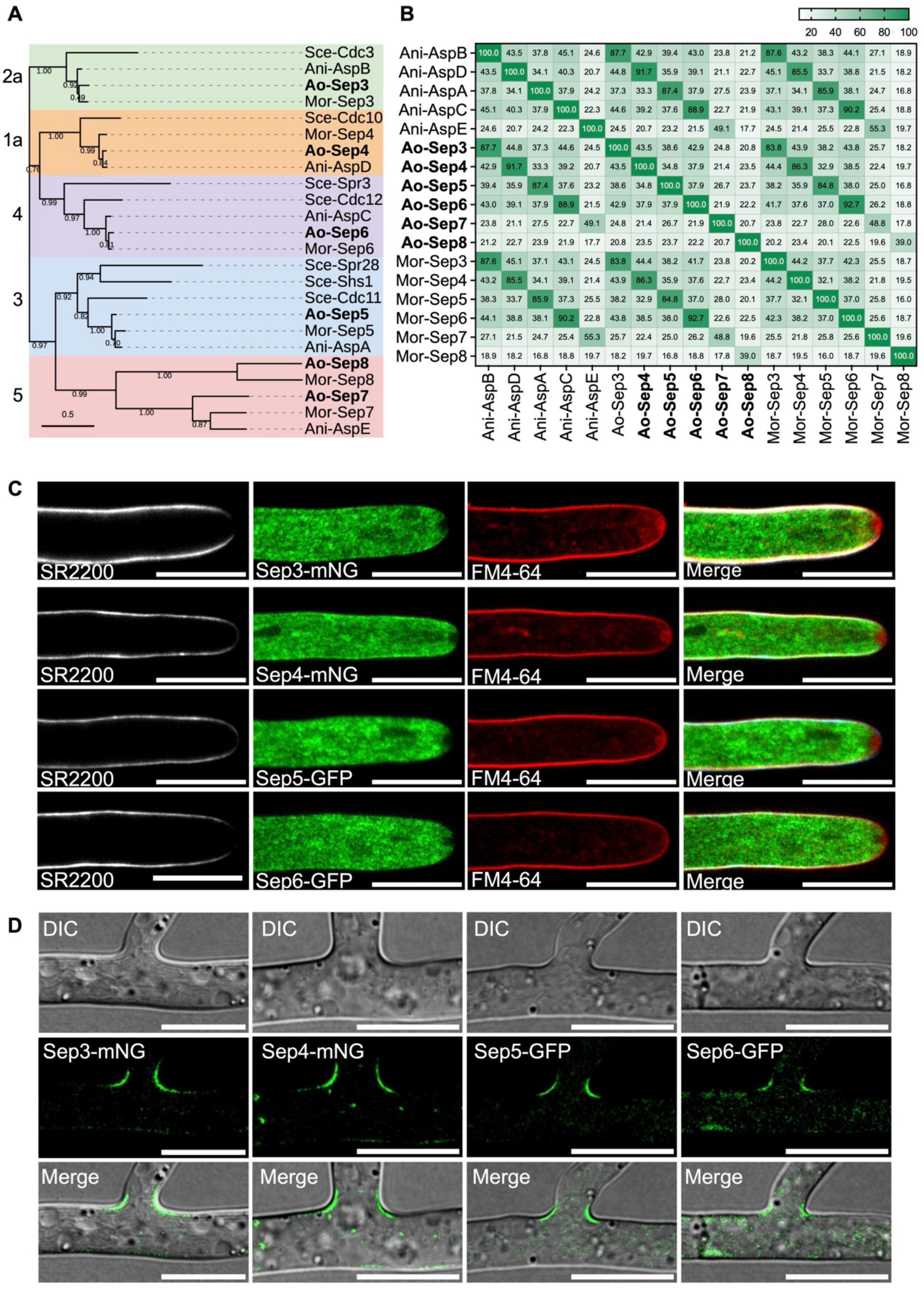
Conserved septins localize at hyphal tips and branch-base curvature in vegetative of *A. oligospora*. **(A)** Phylogenetic analysis of fungal septins. The maximum likelihood tree was inferred using the same approaches as described in Figure 1A. Numbers on the left denote the septin classes. All species abbreviations are consistent with those previously defined. **(B)** Heatmap displaying pairwise amino acid sequence identity among fungal septins. The alignment strategy, color gradient definitions, and numerical descriptors are identical to those described in Figure 1C. Species abbreviations are consistent with those previously defined. **(C)** Localization of Sep3-mNG, Sep4-mNG, Sep5-GFP, and Sep6-GFP in vegetative hyphae. In all panels, hyphae were co-stained with SR2200 to label cell walls and FM4-64 to visualize the plasma membrane and the SPK. Merged images display the SR2200, GFP or mNG, and FM4-64 channels. **(D)** Localization of Sep3-mNG, Sep4-mNG, Sep5-GFP, and Sep6-GFP at hyphal branch points. The merged images show mNG or GFP and DIC channels. All scale bars are 10 μm. Images in all panels are representative of three independent biological replicates.

To examine their localization, we tagged the core septins with GFP or mNG and expressed them under endogenous promoters. In the fungal hyphal tips, we found that the core septins displayed strong diffuse signal in the cytosol but were excluded from the SPK, resulting in a distinct central void within the apical septin distribution (Fig. 2C). They also localized at hyphal branching sites, accumulating at the base of nascent lateral hyphae where the curvature of membrane is pronounced (Fig. 2D). A similar localization pattern has been observed in the hyphae of *A. nidulans* and *Ashbya gossypii* (Bridges et al, 2016; Hernández-Rodríguez et al, 2014), suggesting that the recruitment of septin complexes to regions of high membrane curvature is a conserved feature of filamentous fungal morphogenesis.

### Polarity cues and cytoskeletal dynamics drive trap loop formation

The transition from vegetative growth to a predatory lifestyle in *A. oligospora* is triggered by nematode-derived signals. Within hours of exposure, specialized trap initials emerge, elongate through polarized growth, and subsequently fuse with the parent hypha to generate functional trapping loops (Fig. 3A). To characterize trap development, we tracked the dynamics of chitin synthases, cell-end markers, and cytoskeletons using time-lapse live-cell imaging. At 200 minutes post-exposure (mpe), Chs1-mNG localized prominently to a bright apical punctum corresponding to the SPK of the nascent trap cell and was additionally enriched at newly formed septa (Fig. 3B; Movie EV2). By 230 mpe, Chs1-mNG remained associated with both the SPK and the septa in developing trap cells and became strongly concentrated at the impending hyphal fusion site (Fig. 3B; Movie EV2). At 235 mpe, signal intensity further increased at the fusion interface, coinciding with cell-cell contact (Fig. 3B; Movie EV2). Chs7-mNG displayed a comparable localization pattern, accumulating at septa and converging at the fusion site (Fig. EV3A; Movie EV3). These patterns indicate that multiple CHSs are recruited to active sites of cell wall remodeling including the apical SPK, septa, and the fusion interface throughout trap morphogenesis.

**Figure 3.**
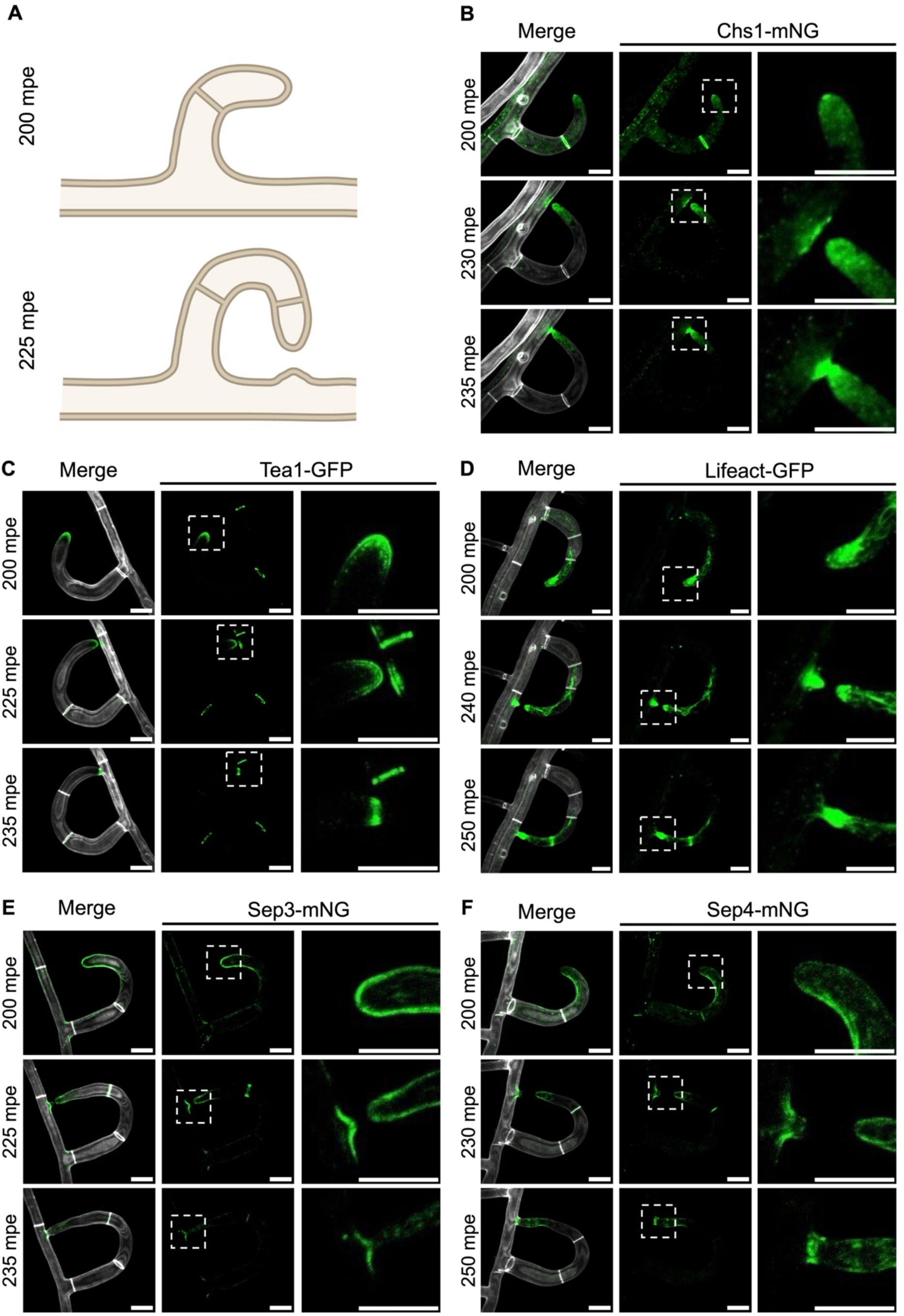
Polarity cues are targeted to fusion sites and cytoskeletal elements are enriched at the inner loop rim during trap formation. **(A)** Schematic representation of the process of trap formation. **(B-F)** Localization of Chs1–mNG **(B)**, Tea1-GFP **(C)**, Lifeact-GFP **(D)**, Sep3-mNG **(E)**, Sep4-mNG, **(F)** during trap formation after a 3-hour exposure to nematodes. Merged panels display the mNG or GFP signals overlaid with SR2200 staining to visualize the cell wall structure. The enlarged images are indicated in the lower-magnification images using white dashed rectangles. All scale bars are 10 μm. Images in all panels are representative of three independent biological replicates.

The cell-end marker Tea1-GFP exhibited a distinct spatial distribution, localizing predominantly to the peripheral cortex of the trap cell apex and at septa (Fig. 3C; Movie EV4). As loop formation progressed, Tea1-GFP accumulated at the prospective fusion site on the vegetative hypha by 225 mpe and remained enriched at this interface through 235 mpe (Fig. 3C; Movie EV4). These dynamics indicate that Tea1 delineates sites of polarized growth and serves as a pre-contact landmark for the fusion interface during trap loop formation.

The actin and septin cytoskeletons, by contrast, displayed distinct temporal dynamics. Lifeact-GFP was enriched at trap hyphal tips and concentrated strongly at the fusion interface by 240 mpe, coincident with the emergence of a vegetative hyphal protuberance (Fig. 3D; Movie EV5). Similarly, Sep3, Sep4, and Sep5 accumulated at branching and fusion regions between 230 and 240 mpe (Fig. 3E and F; Fig. EV3B; Movie EV6-8), whereas Sep6-GFP localized predominantly to developing septa (Fig. EV3C; Movie EV9). Within the expanding loop, both actin and septins were preferentially enriched along the inner concave rim of the trap (Fig. 3D-F; Movie EV5-8). This distribution is consistent with the preferential association of actin and septins with regions of high membrane curvature reported in other system (Bridges et al, 2016; Lou et al, 2019), suggesting that their recruitment is coupled to the specialized membrane geometry of trapping loop formation and expansion in *A. oligospora*.

### Polarity cues and cytoskeletal dynamics mark the site of invasive hyphal emergence within the infection bulb

Following nematode capture, trap cells differentiate infection structures that penetrate the nematode cuticle (Fig. 4A). To characterize infection structures development, we performed time-lapse live-cell imaging to track the dynamics of chitin synthases, cell-end markers, and the cytoskeleton after adding nematodes to pre-formed traps. Developmental progression was monitored over time and is expressed as minutes post-capture (mpc). From 35 to 45 mpc, Chs1-mNG was broadly distributed throughout the developing infection bulb (Fig. 4B; Movie EV10). As morphogenesis shifted toward invasive growth by 75 mpc, Chs1-mNG became intensely enriched at the apical tips of emerging invasive hyphae (Fig. 4B; Movie EV10). The diffuse distribution of Chs1 in the expanding infection bulb likely provides a reservoir for chitin synthesis during subsequent polarized growth of invasive hyphae.

**Figure 4.**
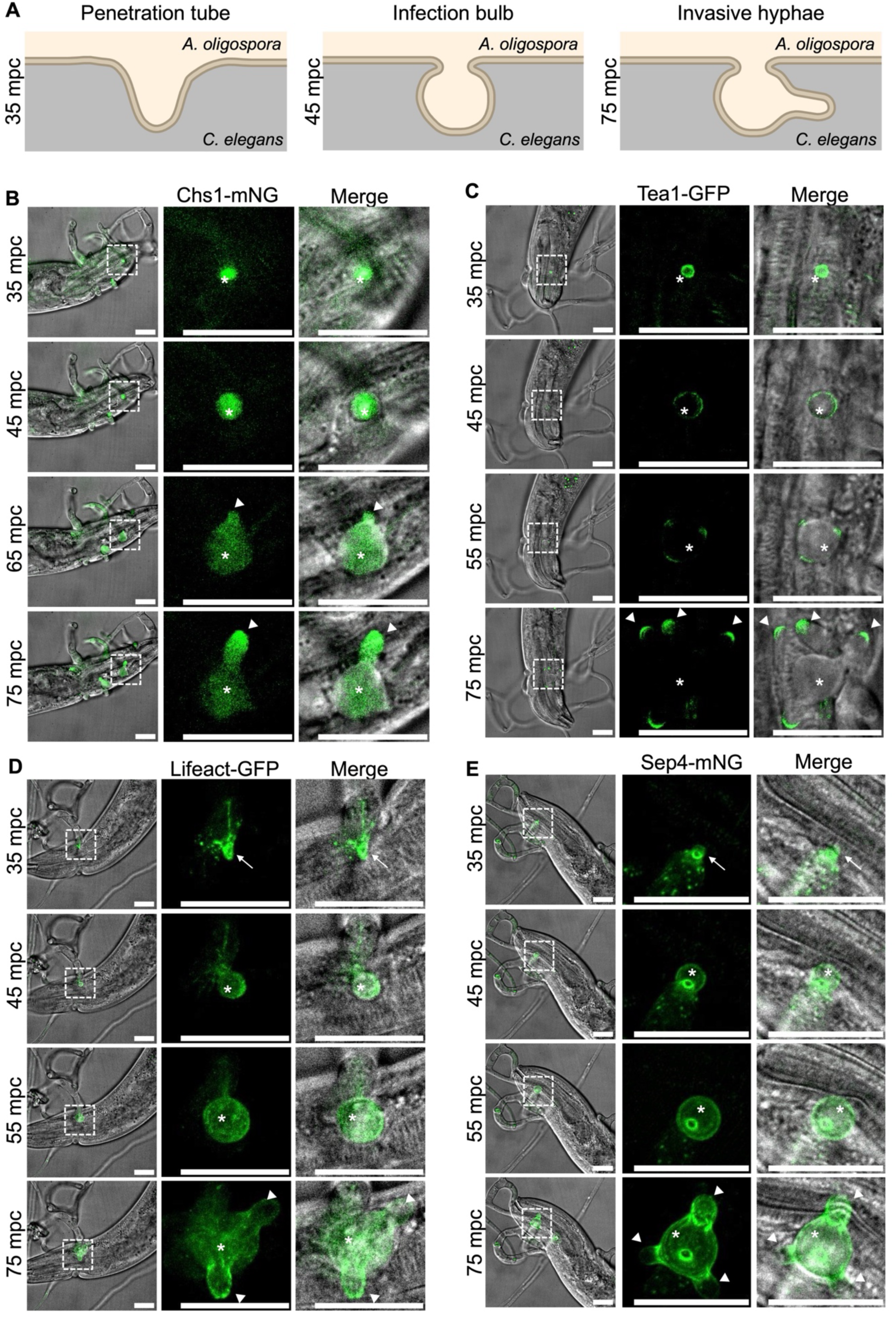
Spatiotemporal localization of polarity cues and cytoskeleton during infection bulb and invasive hypha formation. **(A)** Schematic representation of infection structures differentiated from trap cells during nematode penetration and colonization. **(B-E)** Localization of Chs1–mNG **(B)**, Tea1-GFP **(C)**, Lifeact-GFP **(D)**, and Sep4-mNG **(E)** during infection bulb and invasive hyphae formation. Imaging monitors the interaction between *C. elegans* and pre-formed traps from 35 to 75 minutes post capture (mpc). The penetration tube, indicated by arrows, has penetrated the *C. elegans* cuticle. Subsequently, the infection bulb, indicated by asterisks, is distinguishable as a swollen structure by 45 mpc, while invasive hyphae, indicated by triangles, are observed emerging from the bulb by 75 mpc. Merged panels display the mNG or GFP signals overlaid with DIC channels, which reveal the location of the trapped nematodes. Enlarged images are indicated in the lower magnification images using white dashed rectangles. All scale bars are 20 μm. Images in all panels are representative of three independent biological replicates.

Tea1-GFP exhibited a similar spatiotemporal shift, initially forming a peripheral cortical signal around the infection bulb between 35 and 45 mpc, and then became concentrated at discrete sites of invasive hyphal emergence by 55 mpc (Fig. 4C; Movie EV11). By 75 mpc, Tea1-GFP was concentrated at the tips of the emerging invasive hyphae. Together, these localization dynamics indicate that Tea1 serves as a spatial landmark for invasive hyphal differentiation within the isotropic infection bulb.

During infection bulb maturation, the actin and septin cytoskeletons underwent a coordinated reorganization. By 35 mpc, Lifeact-GFP and core septins assembled into prominent ring-like structures at sites that subsequently developed into infection bulbs (Fig. 4D and E; Fig. EV4; Movie EV12-16). As the infection bulb expanded, both signals became diffuse within the bulb region. However, their localization diverged following invasive hyphal emergence. By 70 mpc, actin accumulated at the tips of the emerging invasive hyphae (Movie EV12). In contrast, septins re-concentrated at the branching point between the infection bulb and the invasive hyphae by 75 mpc (Fig. 4E; Fig. EV4; Movie EV13-16). The divergent recruitment of actin to the hyphal apices and of septins to the branching point reveals the distinct spatial organization of the active hyphal tip and the basal branch site.

### Chitin synthase and cell-end markers coordinate trap morphogenesis

To define the functional contributions of Chs1 and the cell-end markers Tea1 and Tea5, we generated target deletions and confirmed the genotypes via PCR and Southern blotting (Fig. EV5). Both *chs1* and *tea1* mutants exhibited reduced vegetative growth on nutrient-rich medium (PDA), although conidiation remained comparable to the wild type (Fig. 5A). Unlike the zigzag hyphae characteristic of *tea* mutants in *A. nidulans* (Takeshita et al, 2008), the hyphal tips of **tea1** and **tea5** mutants remained straight in *A. oligospora* (Fig. 5B), Notably, the *tea5* mutant developed zigzag hyphae specifically within the colony interior (Fig. 5B), suggesting that Tea5 may primarily govern hyphal network architecture rather than apical growth in *A. oligospora*.

**Figure 5.**
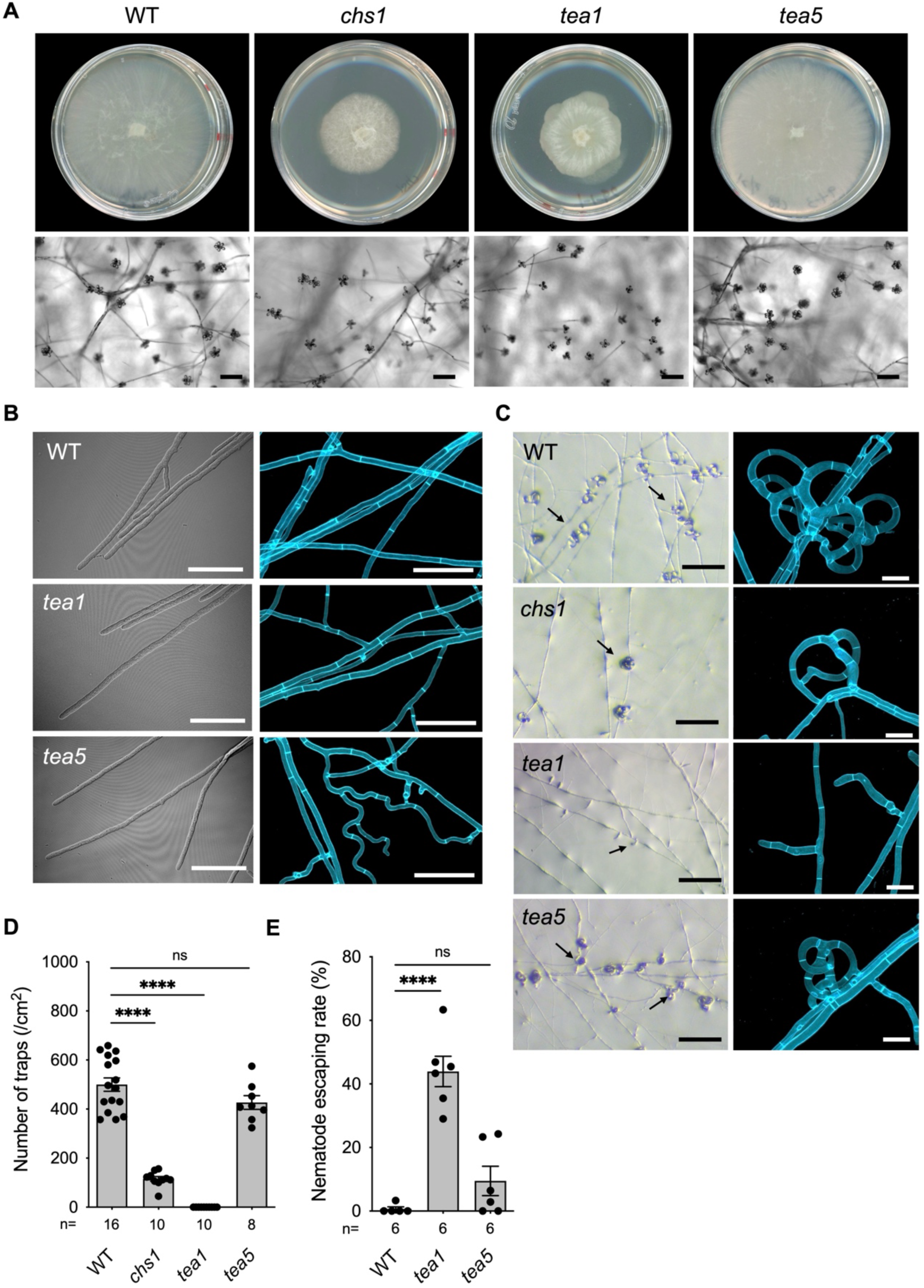
Chitin synthase Chs1 and cell-end markers Tea1 and Tea5 play distinct roles in hyphal growth and trap morphogenesis in *A. oligospora*. **(A)** Colony morphology and conidiation. Upper: colonies of the wild type (WT), *chs1*, *tea1*, and *tea5* mutants grown on 5.5-cm PDA plates for 3 days. Lower: micrographs showing conidiation profiles after culturing for 4 days on PDA. Scale bar: 100 μm. **(B)** Left: Representative DIC images of the wild type and *tea* mutants’ hyphal tips. Right: WT, *tea1* and *tea5* mutants were stained with SR2200 to visualize the interconnected hyphal networks. Scale bar: 50 μm. **(C)** Left: Representative bright field images of the traps induced by *C. elegans* in the WT, *chs1*, *tea1*, and *tea5* mutants. Scale bar: 200 μm. The black arrows indicate traps. Right: Images of traps formed by the WT, *chs1*, *tea1*, and *tea5* strains after 24 hours of continuous induction with 400 *C. elegans larvae*. Confocal images of traps stained with SR2200 after 24 hours of continuous induction with 400 *C. elegans larvae*. Scale bar: 20 μm, respectively. The images are representative of three independent biological replicates. **(D)** Quantification of the trap numbers induced by *C. elegans* in the WT, *chs1*, *tea1*, and *tea5* mutants. **(E)** Quantification of nematode escaping rates following interaction with the WT, *tea1*, and *tea5* mutants. For **(D)** and **(E)**, data represent mean ± SEM; n shown along the x axis; two-tailed unpaired Student’s *t*-test. **** *p* < 0.001; ns, not significant. Images in all panels are representative of three independent biological replicates.

To assess the role of Chs1 and Tea1 in trap morphogenesis, we quantified trap formation after exposure to 30 *C. elegans* L4 stage larvae. The *chs1* mutant developed approximately 100 traps per cm², representing a significant reduction compared to the ∼500 traps per cm² in the wild type (Fig. 5C and D). These *chs1* traps were morphologically defective, typically consisting of only simple hyphal loops (Fig. 5C). The *tea1* mutant exhibited a more severe morphogenetic failure, developing stalk-like initials that lacked the curvature required for loop formation (Fig. 5C and D). In contrast, the *tea5* mutant developed traps in numbers comparable to the wild type but reduced in size (Fig. 5C and D). While both the *tea1* and *tea5* mutants remained capable of capturing nematodes, their predatory efficiency was significantly compromised (Fig. 5E). Together, these findings demonstrate that Tea1 is essential for the directed hyphal bending that drives trap loop formation, whereas Tea5 modulates trap scale and the integrity of the vegetative hyphal network.

### Septins differentially regulate vegetative growth and trap morphogenesis

To investigate the roles of septins in *A. oligospora*, we generated deletion mutants of the four core septins and validated the genotypes by PCR and Southern blotting (Fig. EV5). Functional analysis revealed varying degrees of importance for vegetative growth and asexual development. Both *sep3* and *sep6* mutants exhibited severe growth defects on PDA and significantly attenuated conidiation (Fig. 6A). In contrast, the *sep5* mutant displayed mild growth defects, while the *sep4* mutant developed comparably to the wild type (Fig. 6A). Reintroduction of *SEP3* and *SEP6* into the respective mutant background restored wild-type phenotype, confirming that these defects were specific to the loss of these septins (Fig. EV6A).

**Figure 6.**
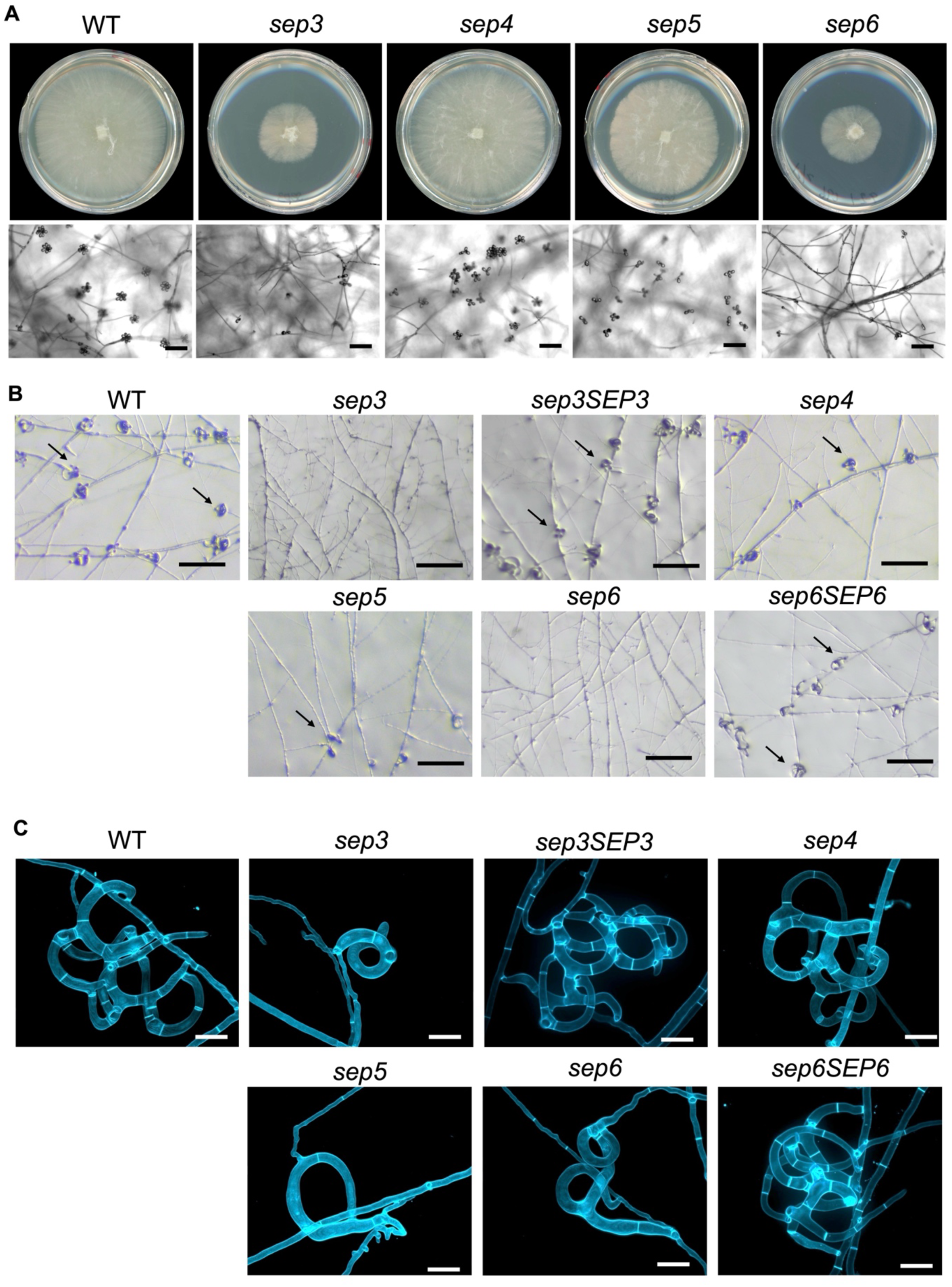
Core septins differentially regulate vegetative growth, conidiation and trap morphogenesis in *A. oligospora*. **(A)** Colony morphology and conidiation. Upper: colonies of the wild type (WT) and *sep* mutants grown on 5.5-cm PDA plates for 3 days. Lower: Micrographs showing conidiation profiles after culturing for 4 days on PDA plates. Scale bars: 100 μm. **(B)** Representative brightfield images of the traps induced by *C. elegans* in the WT and in *sep* mutants, and the *sep3* and *sep6* complementation lines. Scale bars: 200 μm. Black arrows indicate traps. **(C)** Confocal images of traps formed by the WT, *sep* mutants, and rescue strains stained with SR2200 after 24 hours of continuous induction with 400 *C. elegans* larvae. Scale bars: 20 μm. Images in all panels are representative of three independent biological replicates.

We next investigated the role of septins in trap development. In response to *C. elegans,* deletion of any core septin significantly reduced trap formation, with *sep3* and *sep6* producing almost no traps (Fig. 6B; Fig. EV6B). When traps were observed, the *sep3*, *sep5*, and *sep6* mutants predominantly produced simplified structures consisting of single or incomplete loops, while the *sep4* mutant produced traps that were largely comparable in morphology to the wild type (Fig. 6C). Complementation of the respective mutants with *SEP3* and *SEP6* restored both trap abundance and architecture (Fig. 6B and C; Fig. EV6B). Together, these results indicate that core septins broadly promote trap induction, while Sep3, Sep5, and Sep6 are specifically required for the maturation of complex, multi-loop trap structures.

### Nox1–mediated ROS provides a spatial cue for fusion machinery recruitment during trap hyphal fusion

Hyphal fusion is a critical requirement for the closure of trap loop. In various fungi, NADPH oxidase 1 (Nox1) and its regulator NoxR1 regulate hyphal fusion during vegetative growth (Dirschnabel et al, 2014; Nordzieke et al, 2019; Roca et al, 2012), although their role in trap development in NTF remains unclear. Phylogenetic analysis confirmed that the *A. oligospora* genome encodes conserved orthologs of both Nox1 and NoxR1 (Fig. 7A and B). Domain architecture analysis further revealed that Ao-Nox1 contains a characteristic ferric reductase transmembrane component like domain, a FAD-binding 8 domain, and a ferric reductase NAD-binding domain (Fig. 7C). Its regulator, Ao-NoxR1, possesses tetratricopeptide repeat (TPR) motifs and a Phox and Bem1 domain (Fig. 7C). These domains are evolutionarily conserved from fungi to mammals and are required for NoxR recruitment to the plasma membrane to activate Nox (Koga et al, 1999; Takemoto et al, 2011).

**Figure 7.**
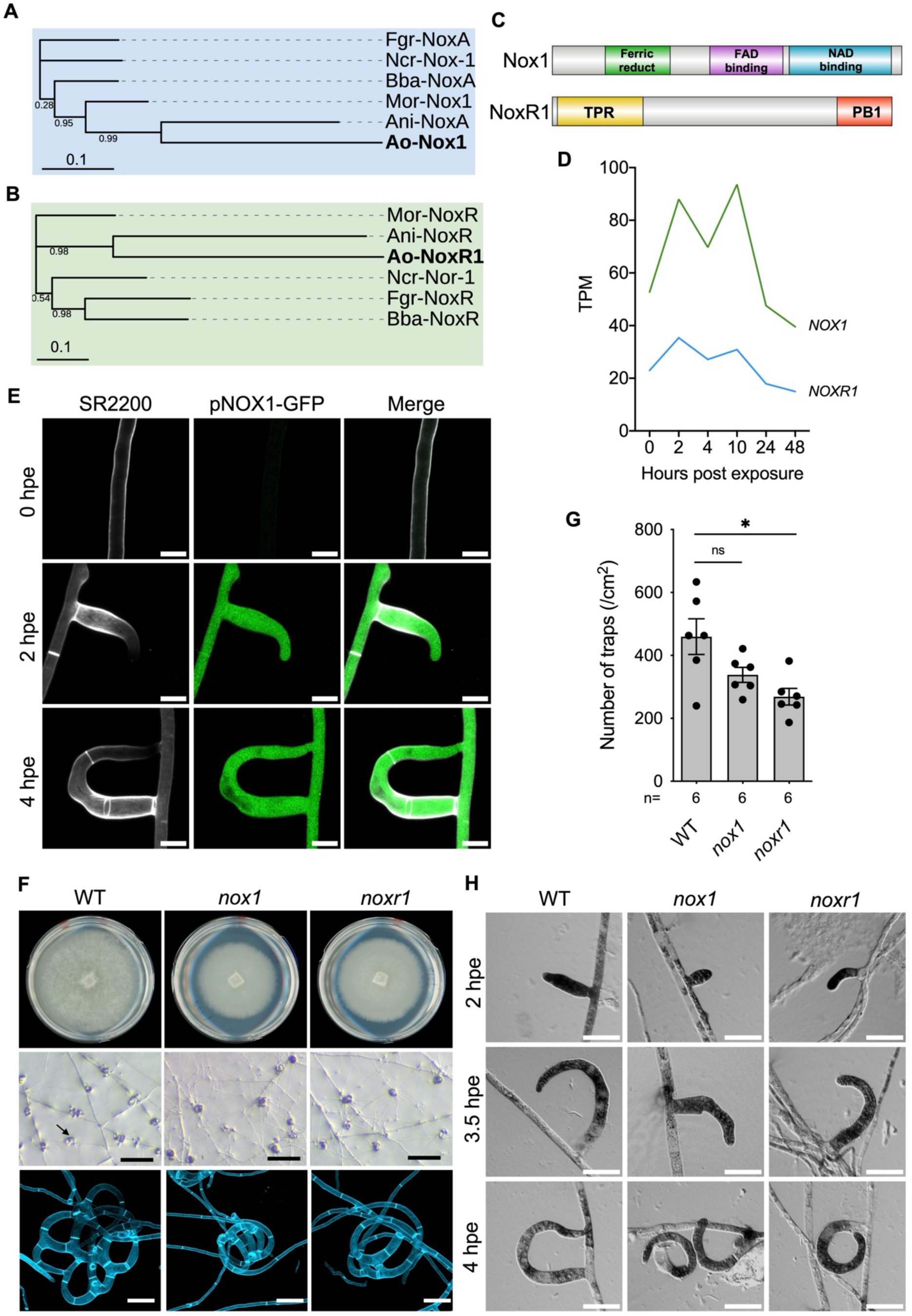
Nox1 and NoxR1 are nematode-responsive and mediate ROS-dependent cell fusion during trap morphogenesis in *A. oligospora*. (A-B) Phylogenetic analysis of fungal Nox1 and NoxR1. Maximum-likelihood tree for Nox1 **(A)** and NoxR1 **(B)** were inferred using FastTree. Node numbers indicate local support values ranging from 0 to 1. The scale bar represents 0.1 amino acid substitutions per site. Species abbreviations are as follows: Ani, *A. nidulans*; Ao, *A. oligospora*; Bba: *Beauveria bassiana*; Fgr, *F. graminearum*; Mor, *M. oryzae*; Ncr, *N. crassa*. **(C)** Domain architecture of Nox1 and NoxR1 in *A. oligospora*. Nox1 proteins contain a ferric reductase transmembrane component-like domain (IPR013130), a FAD-binding 8 domain (IPR013112), and a ferric reductase NAD-binding domain (IPR013121); NoxR1 contains tetratricopeptide repeat (TPR) (IPR011990) and a Phox and Bem1 (PB1) (IPR000270) domain. **(D)** TPM values of the differentially expressed *NOX1* and *NOXR1* in a published RNA-seq time course following exposure of *A. oligospora* to *C. elegans* (Lin et al, 2023). Values represent the average of three independent biological replicates. **(E)** The localization of pNOX1-GFP is displayed at 0, 2, and 4 hours after *C. elegans* exposure. The merged images show GFP and SR2200 channels. Scale bars: 10 μm. Images are representative of three independent biological replicates. **(F)** Upper: colonies of the wild type (WT) and *nox* mutants grown on 5.5 cm PDA plates for 3 days. Middle: Representative brightfield images of the traps induced by *C. elegans* in the WT and in *nox* mutants. Scale bars: 200 μm. Black arrows indicate traps. Lower: Confocal images of traps formed by the WT and *nox* strains stained with SR2200 after 24 hours of continuous induction with 400 *C. elegans* larvae. Scale bar: 20 μm. **(G)** Quantification of the trap numbers induced by *C. elegans* for the WT and *nox* strains. Data represent mean ± SEM; n shown along the x axis; two-tailed unpaired Student’s *t*-test; * *p* < 0.05; ns, not significant. **(H)** ROS production visualized by NBT staining in the WT and *nox* mutants is displayed at 2, 3.5 and 4 hours after *C. elegans* induction. Scale bar: 20 μm. Images are representative of three independent biological replicates. Images are representative of three independent biological replicates.

Transcriptional profiling following nematode exposure revealed that both *NOX1* and *NOXR1* are nematode responsive genes, with transcript levels significantly up-regulated during the initiation of trap development (Fig. 7D). To verify this induction at the cellular level, we monitored a GFP reporter driven by the endogenous *NOX1* promoter. While pNOX1–GFP was undetectable during vegetative growth, the signal was robustly induced specifically within the trap initials upon nematode contact (Fig. 7E). By 4 hpe, the GFP signal remained intensely concentrated throughout the elongating and curving trap cells (Fig. 7E). This highly specific expression pattern demonstrates that the activation of the Nox1 complex is strictly coupled with the onset of trap morphogenesis and suggests its involvement in the specialized cellular processes required for predation.

To characterize the function of NOX enzymes in trap development, we generated *nox1* and *noxr1* deletion mutants. Deletion of *NOX1* and *NOXR1* caused mild growth defects but severe trap morphology abnormalities (Fig. 7F). Upon exposure to *C. elegans*, the *nox* mutants produced trap numbers comparable to the wild type; however, these structures consistently failed to undergo hyphal fusion (Fig. 7F and G). Instead of arresting growth and integrating into the parent hypha to close the loop, the mutant hyphae bypassed the prospective fusion site, resulting in characteristic pigtail-shaped structures (Fig. 7F). This phenotype demonstrates that Nox1 and NoxR1 are dispensable for trap initiation and elongation but are essential for the site-specific fusion event required for loop closure.

To determine if this fusion failure stems from defective signaling, we monitored ROS production using nitroblue tetrazolium (NBT) staining to detect superoxide radicals generated by NOX enzymes. During early trap development (2–3.5 hpe), superoxide accumulation was detected at the hyphal tips and nascent traps of both wild-type and *nox* mutant strains, indicating that early ROS production is likely Nox1-independent (Fig. 7H). Notably, by 4 hpe, a localized NBT signals appeared at the specific fusion sites on the receiving hyphae in the wild type, whereas this signal was absent in the *nox1* and *noxr1* mutants (Fig. 7H). These results indicate that Nox1 and NoxR1 are specifically required to generate a localized ROS signal on the target hypha to trigger cell-cell fusion during trap development.

We finally investigated whether Nox1-mediated ROS signaling is required to recruit the cell polarity proteins and cytoskeletal machinery to the fusion interface. In the wild type, Chs1-mNG, Tea1-GFP, and Lifeact-GFP accumulated intensely at both the trap tips and corresponding fusion sites on the receiving hyphae by 225 mpe (Fig. 8 A Movie EV17-19). In contrast, while these proteins localized normally to the trap tips in the *nox* mutants, they failed to accumulate at the prospective fusion site on receiving vegetative hyphae (Fig. 8B and C; Movie EV17-19). Consequently, by 300 mpe, the mutant traps remained unfused and retained their aberrant pigtail morphology (Fig. 8B and C; Movie EV17-19). Together, these findings demonstrate that Nox1– mediated ROS production serves as a critical spatial signal required for the recruitment of the fusion machinery, ensuring trap loop closure and the integration of the predatory network.

**Figure 8.**
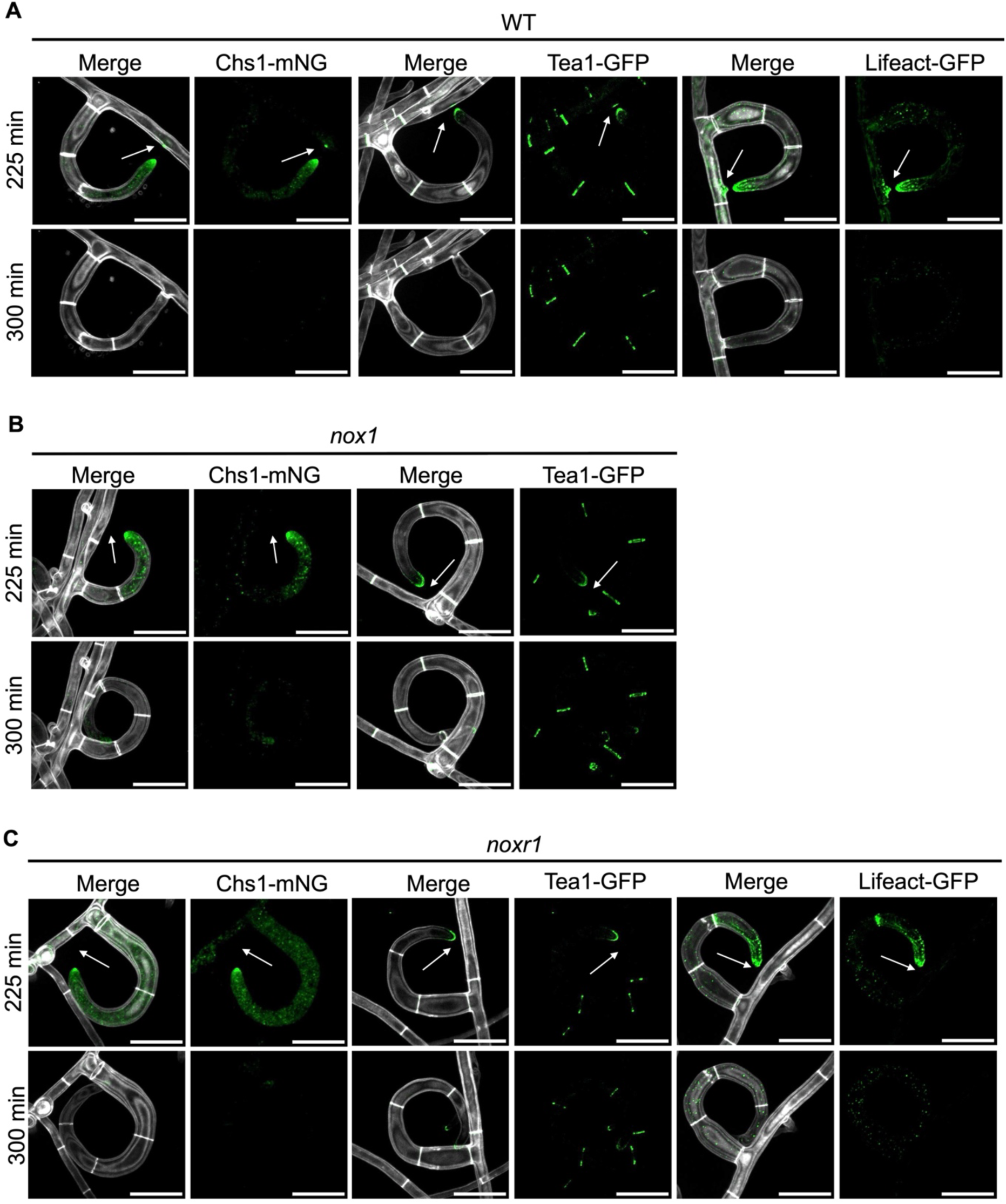
Nox1 and NoxR1 are required for recruitment of polarity cues and the actin at fusion sites during trap formation. (A-C) Localization of Chs1-mNG, Tea1-GFP, and Lifeact-GFP in the wild type (WT) **(A)**, *nox1* **(B)**, and *noxr1* **(C)** during trap formation at 225 to 300-minutes after exposure to nematodes. Merged panels display the mNG or GFP signals overlaid with SR2200 channels, which reveal the cell wall structure. All scale bars are 20 μm. White arrows indicate prospective fusion sites. Images in all panels are representative of three independent biological replicates.

## Discussion

The morphogenesis of nematode-trapping structures represents a striking example of cellular reprogramming, where conserved growth modules are redeployed to generate complex three-dimensional architectures. Here, we elucidate the cellular and molecular mechanisms underlying this transition in *A. oligospora*, revealing how cell polarity proteins, cytoskeletal mechanics, and Nox1-mediated ROS signaling are coordinately integrated to build a functional predatory apparatus (Fig. 9A).

**Figure 9.**
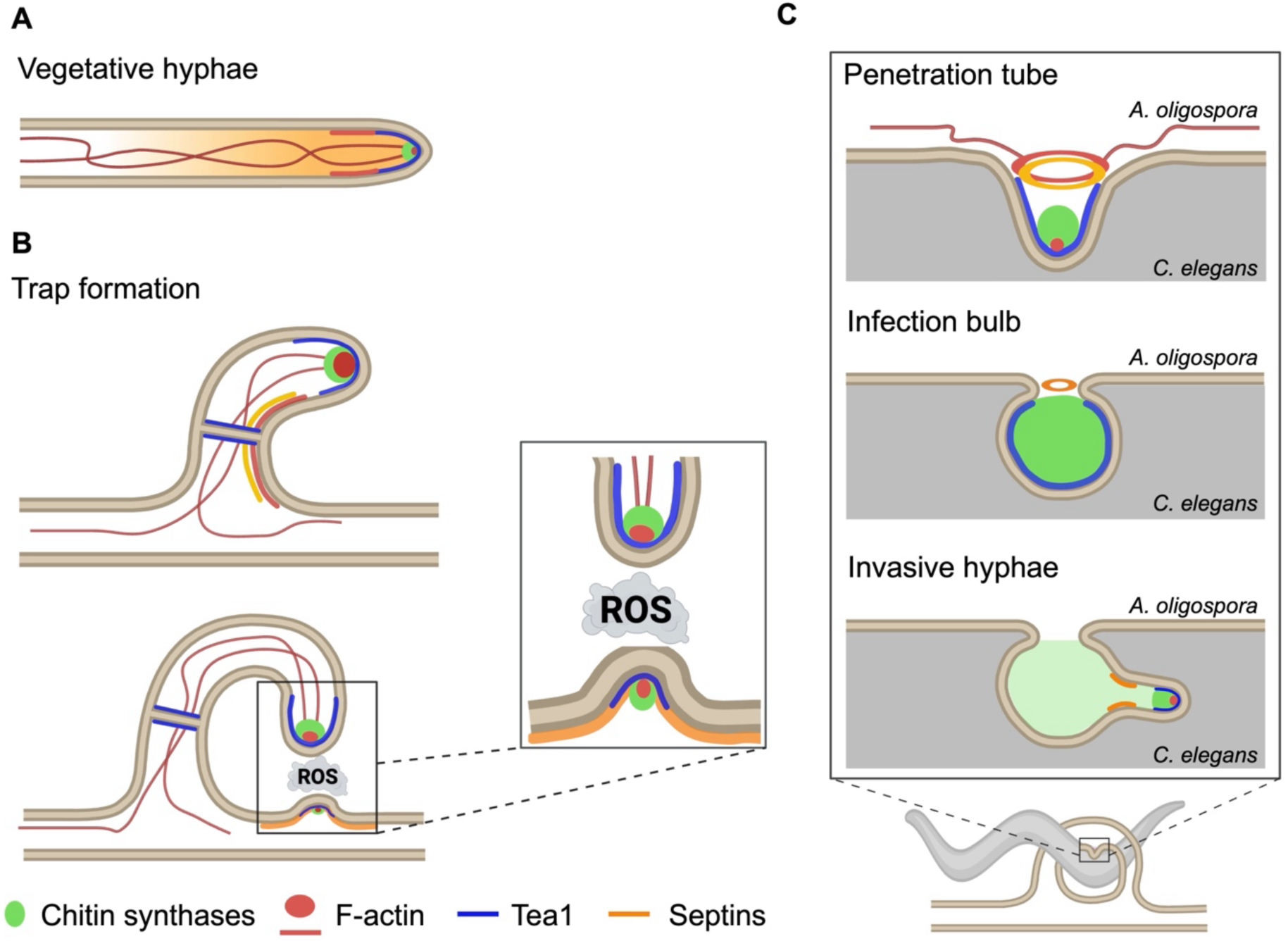
Hypothetical model of cell polarity cues, cytoskeletal dynamics, and NADPH oxidase signaling across the predatory lifecycle of *A. oligospora*. **(A)** During polarized hyphal growth, the chitin synthases Chs1 and Chs7 (green) localize to the spitzenkörper (SPK) and hyphal tips, respectively, for cell wall synthesis. F-actin (red) accumulates at the tip and in subapical patches. Tea1 (blue) marks the plasma membrane at the hyphal apex, where it is proposed to serve as a positional cue for growth orientation. Septins (orange) form a diffuse structure at the hyphal tip while being excluded from the SPK. **(B)** Upon nematode detection, trap cells emerge from vegetative hyphae and undergo curved growth. Chitin synthases, Tea1, F-actin, and septins relocalize to trap cell tips during curvature formation. F-actin and septins show enrichment along the inner rim of developing loops, contributing to trap curvature. At the fusion site (lower panel), Nox1-mediated ROS production (indicated by cloud) is essential for recruiting chitin synthases, Tea1, F-actin, and septins in the vegetative hyphae to enable cell fusion and complete loop closure, forming functional trapping structures. Additionally, septins localize to the branch points at the fusion site where new hyphae emerge. **(C)** After trap capture of nematodes, infection structures develop. Initially, a penetration tube forms to breach the nematode cuticle (upper panel). Subsequently, infection bulbs differentiate within the nematode body (middle panel). F-actin and septins assemble into ring-like structures at sites where infection bulbs form, and Tea1 accumulates around the bulb, while chitin synthases become diffusely distributed throughout the bulb. Later, chitin synthases and Tea1 concentrate at invasive hyphal tips emerging from infection bulbs (lower panel), directing penetration and colonization of the nematode host.

The transition from vegetative growth to trap formation requires a spatial shift in the polarity machinery (Fig. 9A and B). While Tea1 and Chs1 exhibit canonical apical localization to support linear extension during vegetative growth (Fig. 9A), they are repurposed to navigate the complex curvature of loop formation upon nematode induction (Fig. 9B). The *tea1* mutant produces straight, stalk-like structures, which indicates that Tea1 specifies the spatial coordinates required for the directed hyphal bending rather than simply promoting linearity as described in *S. pombe* and *A. nidulans* (Mata & Nurse, 1997; Takeshita et al, 2013). Notably, the persistence of Tea1 at septa in *A. oligospora* suggests that polarity markers are retained following cytokinesis and may be rapidly redeployed during trap development. Similar observations in *A. flagrans (Kriegler et al, 2025b)* raise the possibility that septum-associated polarity retention is a conserved feature among NTF, although the underlying regulatory mechanisms remain unknown. In contrast, Tea5 regulates hyphal network organization and trap size in *A. oligospora*, with phenotypes distinct from those reported for orthologs in *A. nidulans (Takeshita et al, 2008). Together,* these observations highlight functional diversification of conserved cell-end markers in NTF to support predation-specific morphogenetic programs.

In parallel with apical extension, the actin and septin cytoskeletons establish a specialized cortical domain along the inner rim that may promote hyphal curvature. During vegetative growth, septins in *A. oligospora* form apical domes and septation rings, consistent with observations in *C. cinerea* (Kakizaki et al, 2023) (Fig. 9A). During trap formation, both actin and septins become transiently enriched along the concave inner rim before dissipating as elongation proceeds (Fig. 9B). This localization is consistent with their established roles in sensing and stabilizing membrane curvature in other eukaryotic systems (Bridges et al, 2016; Kessels & Qualmann, 2021). We propose that actin and septins function as a transient mechanical scaffold that stabilizes curved membranes during rapid loop expansion and fusion. Given that septins bind specific phosphoinositides or sphingolipids in other fungi (Casamayor & Snyder, 2003; Mela & Momany, 2021), trap induction likely generates localized lipid asymmetries at the inner rim, creating a geometric lipid interface that recruits septins to the concave face. Whether such lipid asymmetry distinguishes the inner and outer rims of the trap is a critical question that needs to be addressed in the future to understand how fungi encode geometric information.

A central feature of our model is the requirement of Nox1-mediated ROS signaling in controlling cell fusion for trap loop closure. The characteristic “pigtail” morphology of *nox1* and *noxr1* mutants effectively uncouples curvature from fusion: while the mutant trap hyphae bend appropriately, they fail to terminate through fusion with the target hyphae. This fusion defect persists even upon physical contact, indicating that Nox1-dependent ROS signaling is essential to recruiting cell polarity cues and cytoskeletal components to the receiving hypha (Fig. 9B, lower panel). Since Nox2-mediated ROS production persists in *nox1* and *noxr1* mutants, we conclude that Nox1 generates a specific localized ROS signal necessary for inter-hyphal communication. We propose that in *A. oligospora*, this ROS burst serves as a fusion checkpoint to ensure that trap loops are fully sealed before predation begins. This mechanism is likely integrated into the pheromone-responsive MAPK pathway, which we previously showed regulates *NOX* transcription (Chen et al, 2021), mirroring the fusion-control pathways (Cano-Domínguez et al, 2008; Leeder et al, 2013).

Following nematode capture, polarity and cytoskeletal modules are reorganized again to drive host penetration (Fig. 9C). The septin and F-actin ring-like structures that we observed at sites of infection bulb formation resemble the septin collars in the appressoria of plant pathogens like *M. oryzae* (Dagdas et al, 2012). Tea1 accumulation in *A. oligospora* during infection bulb development mirrors its role in *M. oryzae*, where it mediates cell polarity to organize the appressorium required for plant infection (Qu et al, 2022). Subsequently, enrichment of CHSs and Tea1 at *A. oligospora* invasive hyphal tips underscores the conserved role of polarity cues in directing host penetration. These parallels indicate that distantly related fungi deploy shared polarity and cytoskeletal modules to execute distinct differentiation programs during host invasion. Together, our study demonstrates that *A. oligospora* has rewired conserved fungal modules governing polarity, cytoskeletal organization, and hyphal fusion to meet the unique mechanical demands of a predatory lifestyle. More broadly, the mechanistic parallels between NTF traps and the appressoria of plant pathogens highlight convergent evolutionary solutions for host penetration (Eisermann et al, 2023). These similarities position NTF as a powerful model for dissecting how eukaryotic cells integrate environmental signals to build complex three-dimensional structures.

## Methods

### Strains and culture conditions

Fungal strains were maintained routinely on potato dextrose agar (PDA; Difco). Trap induction and nematode escaping rate assays were all performed on low-nutrient medium agar (LNM: 2% agar, 1.66 mM MgSO_4_, 5.4 µM ZnSO_4_, 2.6 µM MnSO_4_, 18.5 µM FeCl_3_, 13.4 mM KCl, 0.34 µM biotin, and 0.75 µM thiamin). Liquid culture of fungi to acquire protoplasts was conducted in potato dextrose broth (PDB; Difco). *Caenorhabditis elegans* N2 (Bristol) was maintained on standard nematode growth medium with *Escherichia coli* (OP50) as the food source. All fungal strains used are listed in Table EV2.

### Bioinformatic analyses of chitin synthases, cell-end markers, septins, and NOX enzymes in the *A. oligospora* genome

The amino acid sequences of each chitin synthase, cell-end marker homolog, septin family members and NOX enzymes from *Saccharomyces cerevisiae*, *Schizosaccharomyces pombe*, *Aspergillus nidulans*, *Arthrobotrys flagrans*, *Neurospora crassa*, and *Magnaporthe oryzae*, and *Fusarium graminearum*, were used as queries to identify their homologs in *A. oligospora* using local BLAST analysis in Blast2GO 5 PRO (Conesa et al, 2005).

To construct phylogenies of chitin synthases, cell-end markers, septins, and NOX enzymes , multiple sequence alignments of proteins from the same family were generated using the T-Coffee program (Taly et al, 2011). Pairwise sequence identity scores were calculated from the resulting alignment. Phylogenetic trees were constructed using FastTree (Price et al, 2009) and visualized using iTOL (Letunic & Bork, 2007). Protein domain predication for each gene was generated using InterPro (Blum et al, 2024). A hypothetical model of polarized growth and trap development in *A. oligospora* was illustrated in BioRender (https://biorender.com).

### Construction of fluorescent fusion plasmids

A complete list of plasmids used and generated is presented in Table EV2. Fluorescent protein constructs for Lifeact, cell-end markers, chitin synthases, and septins were generated using GFP or mNeonGreen. Briefly, each target gene, along with a 2 kb upstream sequence, was amplified using specific primers (all primer sequences are in Table EV3) and cloned into a plasmid containing a geneticin resistance cassette and encoding the respective fluorescent protein. The resulting plasmids were transformed into TWF154 strain, and transformants carrying fluorescent-protein-tagged target genes were examined using fluorescence microscopy.

### Live cell imaging of fungal development

To assess the localization of chitin synthases, cell-end markers, F-actin, and septins, strains expressing fluorescent protein fusions were inoculated onto 5.5-cm LNM plates and incubated at 25°C. Vegetative hyphal tips were imaged after 2 days of growth, whereas 3-day-old cultures were utilized as the starting material for subsequent trap induction and infection structures differentiation. For vegetative hyphae, cultures were stained with SCRI Renaissance 2200 (SR2200) and FM4-64 to label fungal cell walls and cell membranes, respectively. Imaging was performed using an LSM980 Airyscan 2 confocal microscope (Carl Zeiss) equipped with a Plan-Apochromat 100×/1.46 oil DIC objective. Images were acquired with Multiplex Mode (SR-4Y) and processed using Zeiss ZEN Blue software.

For developmental analysis, *A. oligospora* cultures were exposed to approximately 400 *C. elegans* L4-stage larvae per plate to induce trap formation. To monitor trap development, nematodes were removed after a 3-hour induction period by washing with deionized water; for infection-structure analysis, mycelia were exposed for 24 hours. To capture the initial stages of penetration, nematodes were introduced to established traps and incubated for 10 minutes before proceeding to imaging. Fungal cell walls and plasma membranes were visualized by staining cultures with SR2200 and FM4-64, respectively. All imaging was performed using an Andor Revolution WD system with a Nikon Ti-E automatic microscope and an Andor iXON Ultra 888 EMCCD camera. A CFI Apochromat TIRF 100×/1.49 oil objective was used for visualization.

### Generation of gene deletion mutants and complementation strains

Targeted gene deletions were carried out in a *ku70* mutant strain of *A. oligospora* TWF154, as its inherent NHEJ deficiency boosts the frequency of homologous. In brief, 1.5 kilobases (kb) of the 5’ and 3’ regions flanking the open reading frame (ORF) of the target gene were amplified and fused to a clonNAT resistance cassette (amplified from plasmid pRS41N) or a geneticin resistance cassette (amplified from plasmid pUMa1057) to generate the knockout constructs. The constructs were then introduced into protoplasts of the *ku70* background (*A. oligospora* TWF1697) via PEG-mediated transformation. To acquire fresh protoplasts, two 5.5-cm plates of *A. oligospora* TWF1697 cultured on PDA were inoculated in 50 mL of PDB and incubated for 24 h at 25°C at 200 rpm. Fungal hyphae were collected by centrifugation and washed using MN buffer (0.3 M MgSO_4_, 0.3 M NaCl). Blended mycelia were mixed with 50 mg/mL VinoTastePro lytic enzyme in MN buffer overnight at 25°C at 200 rpm. Protoplasts were collected by filtering the mixture through 3 layers of Miracloth and washed with STC buffer (1.2 M sorbitol, 10 mM Tris-HCl pH 7.5, 50 mM CaCl_2_). Next, 5 🞨10^5^ protoplasts were mixed with 3 μg of construct DNA and incubated on ice for 30 min. Five volumes of PTC buffer [40% PEG 4000 (w/v), 10 mM Tris-HCl pH 7.5, 50 mM CaCl_2_) was then added and allowed the mixture was incubated for 20 min at room temperature. Then, molten regeneration agar (1% agar, 0.5 M sucrose, 0.3% yeast extract [w/v], 0.3% casein acid hydrolysate [w/v]) containing 180 μg/mL clonNAT or 220 μg/mL geneticin was added to the protoplasts and poured onto Petri dishes. Transformants were then screened for gene deletion. Primers used for gene deletion and genotyping are listed in Table EV3.

Successful knockouts of target genes were first confirmed by polymerase chain reaction (PCR), and then validated by Southern blotting (Fig. S5) to establish the presence of the ectopically integrated drug cassette as described before (Chen et al, 2021). In transgenes intended for complementation of deletion strains, a copy of the wild-type locus containing 1.5–2 kb upstream and 1 kb downstream of the ORF was fused with a geneticin resistance cassette amplified from plasmid pUMa1057 by PCR and transformed into the protoplasts of targeted mutant strains.

### Phenotypic characterization

Trap numbers formed by *A. oligospora* in response to *C. elegans* were quantified as described previously (Yang et al, 2020). Briefly, fungal strains were grown on 3.5-cm LNM plates at 25°C for 48 hours. Subsequently, thirty *C. elegans* L4 stage larvae were introduced onto the fungal plates and incubated for 6 hours before removal. After 24 hours of nematode exposure, three micrographs within 0.5 cm of the plate edge were captured using a Zeiss Stemi 305 microscope with a Zeiss Axiocam ERc 5s camera at 40× magnification. Trap numbers were quantified from the three images.

To compare trap morphology and quantify the nematode escape rate in wild type and mutant strains, fungi were inoculated onto 5.5-cm LNM plates and incubated at 25°C for 3 days. The cultures were then exposed to approximately 400 *C. elegans* L4-stage N2 larvae per plate for 24 hours to induce trap formation. For trap morphology analysis, cultures were stained with SR2200 to label fungal cell walls. Imaging was performed using an LSM980 Airyscan 2 confocal microscope (Carl Zeiss) equipped with a Plan-Apochromat 40×/1.2 water DIC objective. Images were acquired with Multiplex Mode (SR-4Y) and processed using Zeiss ZEN Blue software. For the nematode escape rate quantification, thirty adult nematodes were placed onto established traps and allowed to crawl on the fungal lawn for 10 min. Untrapped nematodes were removed by gently washing the plate with deionized water. The nematode escape rate was calculated as the proportion of untrapped nematodes relative to the total number tested.

### Detection of superoxide during trap development

Superoxide was detected by staining fungal hyphae with nitroblue tetrazolium (NBT), which is reduced by superoxide to form a dark blue, water-insoluble formazan. Trap induction was performed as described in the previous section. To visualize superoxide, 1 mL of freshly prepared 0.1% (w/v) NBT in 0.05 M sodium phosphate buffer was added to each plate and incubated for 20 min at room temperature in the dark. Excess dye was removed by rinsing twice with PBS, and samples were imaged immediately. Imaging was performed on an LSM980 Airyscan 2 confocal microscope (Carl Zeiss) equipped with a Plan-Apochromat 40×/1.2 water DIC objective. Images were acquired in Multiplex Mode (SR-4Y) and processed with Zeiss ZEN Blue software.

### Statistical analysis

For trap quantification, statistical analyses were performed with data from at least three independent biological replicates using two-tailed unpaired Student’s *t* tests or one-way analysis of variance followed by uncorrected Fisher’s least significant difference multiple test. *P* < 0.05 was considered statistically significant.

## Data availability

The *A. oligospora* TWF154 genome sequence used in this study is from NCBI (accession number SOZJ00000000). RNA-seq data used in this study are from the NCBI GEO (accession number GSE233568). The data that support the findings of this study are available from the corresponding author upon request.

## Acknowledgments

We thank Dr. Norman van Rhijn for providing fluorescent protein constructs. We thank Dr. Yae-Huei Liou, Sue-Ping Lee, and Chun-Yung Chang at the Imaging Core at the Institute of Molecular Biology, Academia Sinica for technical assistance with imaging. We are grateful to Dr. Libera Lo Presti for her comments and suggestions of our manuscript. Funding for this work was provided by the Max Planck Society and Academia Sinica Investigator Award (to Y.-P.H., AS-IA-111-L02).

## Author Contributions Statement

C.-Y.K., and Y.-P.H. conceived and designed the research. C.-Y.K., H.-C.L., Y.-Z.C., and S.-A.C. performed the experiments. C.-Y.K., H.-C.L., Y.-Z.C., and S.-A.C., and Y.-P.H. analyzed the data. C.-Y.K., H. S. and Y.-P.H. wrote the manuscript.

---

**Figure EV1.**
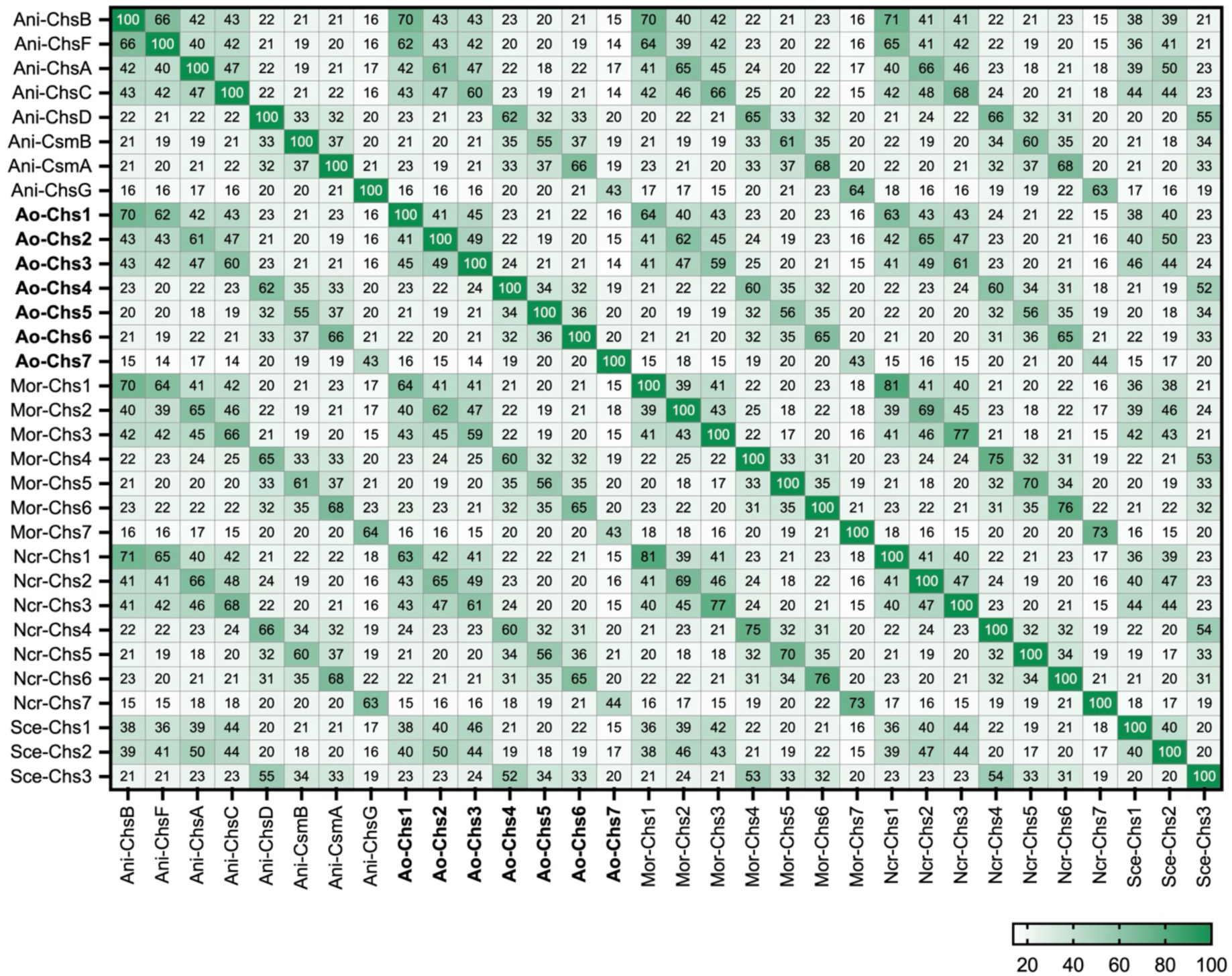
Evolutionary conservation of chitin synthases (CHSs) and nematode-responsive expression of CHS and septin genes. Heatmap displaying pairwise amino acid sequence identity among fungal CHSs. The matrix relies on a multiple sequence alignment generated using the T-Coffee program. The color gradient signifies the degree of conservation, where darker shading corresponds to higher identity percentages. Numerical values within the cells represent percent identity for each protein pair. Species abbreviations are as follows: Ani, *Aspergillus nidulans*; Ao, *Arthrobotrys oligospora*; Mor, *Magnaporthe oryzae*; Ncr, *Neurospora crassa*; Sce, *Saccharomyces cerevisiae*.

**Figure EV2.**
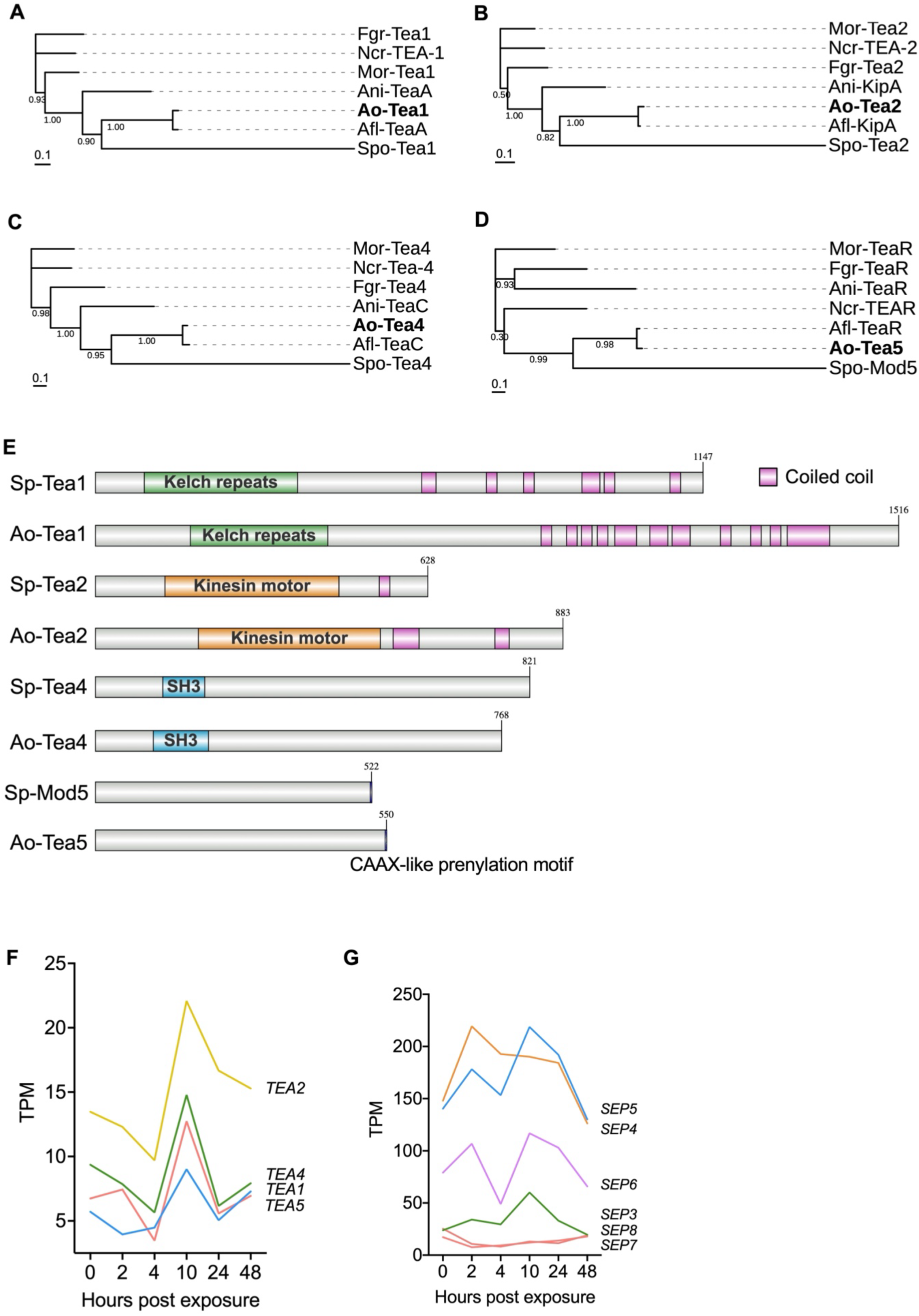
Phylogeny, domain architecture, and expression of cell end markers. (A-D) Phylogenetic analysis of cell end marker proteins. Maximum-likelihood trees for Tea1 **(A)**, Tea2 **(B)**, Tea4 **(C)**, and Tea5 **(D)** were inferred using FastTree. Species abbreviations are as follows: Ani, *A. nidulans*; Ao, *A. oligospora*; Fgr, *Fusarium graminearum*; Mor, *M. oryzae*; Ncr, A. *N. crassa*; Sce, *S. cerevisiae*; Spo, *Schizosaccharomyces pombe*. **(E)** Domain architecture of Tea1, Tea2, Tea4, Mod5, and Tea5 proteins in *S. pombe* and *A. oligospora*. Tea1 proteins in both species contain kelch-repeat domains and C-terminal coiled-coil regions; Tea2 possesses kinesin motor and coiled-coil domains, while Tea4 is characterized by an SH3 domain. Both Mod5 and Tea5 feature a C-terminal CAAX-like prenylation motif. **(F** and **G)** TPM values of the differentially expressed cell-end markers **(F)** and septins **(G)** in a published RNA-seq time course following exposure of *A. oligospora* to *C. elegans* (Lin et al, 2023). Values represent the average of three independent biological replicates.

**Figure EV3.**
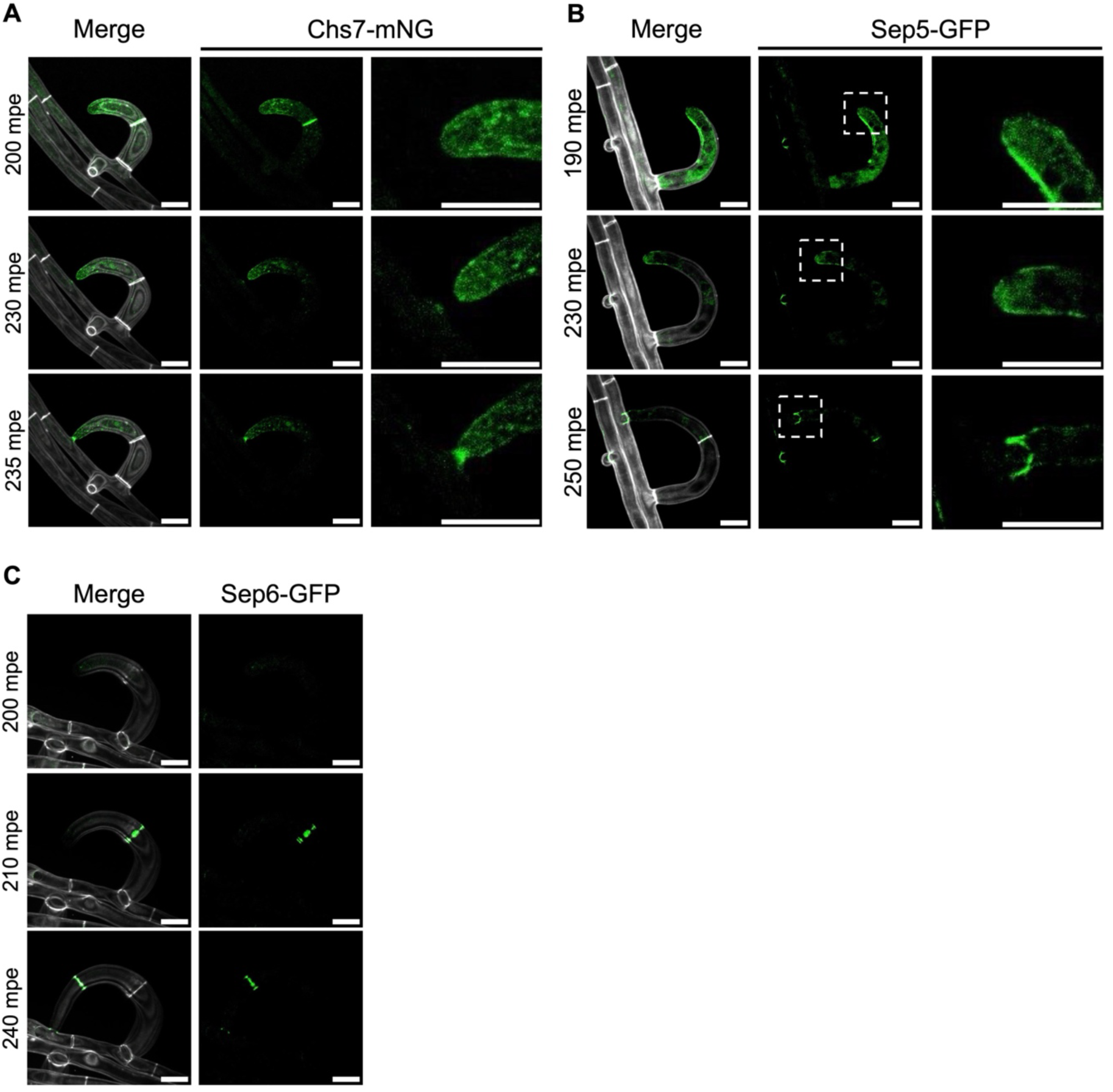
The localization of Chs7-mNG and Sep6-GFP during trap formation in A. oligospora. (A-C) Localization of Chs7–mNG **(A)**, Sep5-GFP **(B),** and Sep6-GFP **(C)** during trap formation at 200-240 minutes after exposure to nematodes. Merged panels display the mNG or GFP signals overlaid with SR2200 channels, which reveal the cell wall structure. The enlarged images are indicated in the lower magnification images using white dashed rectangles. All scale bars are 10 μm. Images in all panels are representative of three independent biological replicates.

**Figure EV4.**
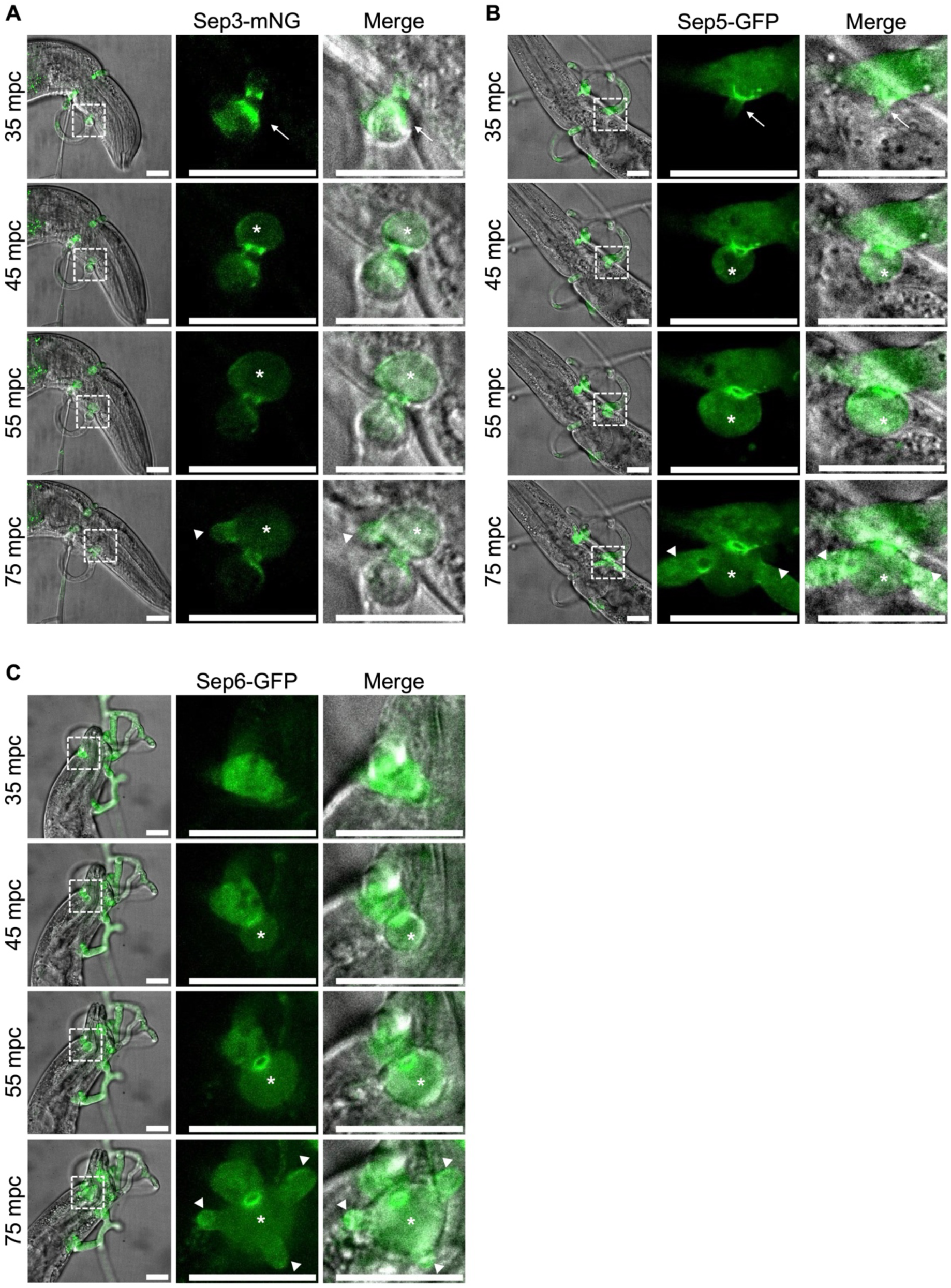
Septins coordinate infection structure differentiation in *A. oligospora*. (A-C) Localization of Sep3–mNG **(A)**, Sep5-GFP **(B)**, and Sep6-GFP **(C)** during infection bulb and invasive hyphae formation at 35-75 minutes after exposure to nematodes. The merged images show mNG or GFP and DIC channels. The enlarged images highlighted by white dashed squares. All scale bars are 20 μm. Images are representative of three independent biological repeats.

**Figure EV5.**
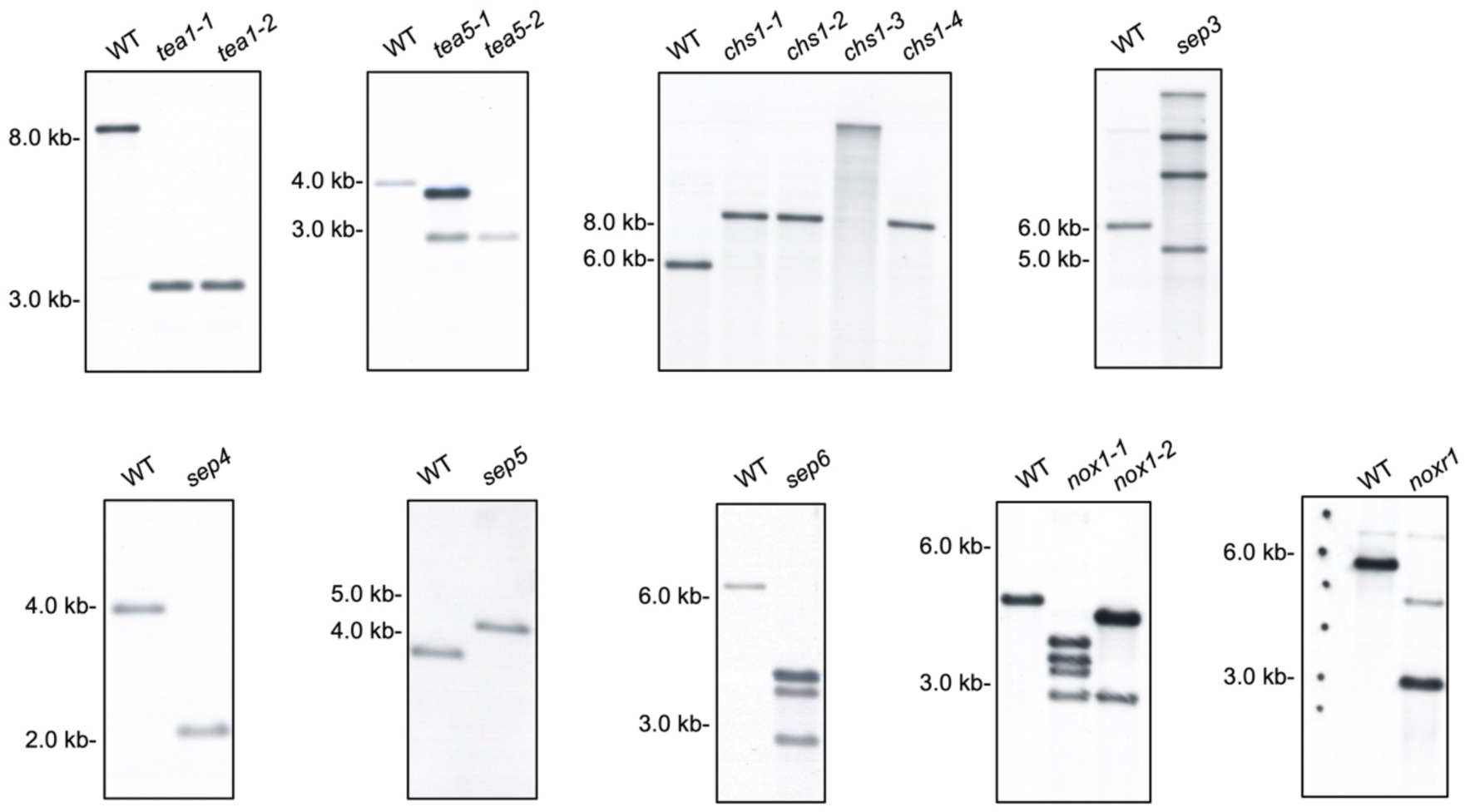
Southern blot analysis of targeted gene deletion mutants. Southern blot analysis was used to confirm the successful deletion of *TEA1*, *TEA5*, *CHS1*, *SEP3*, *SEP4*, *SEP5*, *SEP6*, *NOX1*, and *NOXR1* via homologous recombination. Genomic DNA from the wild type (WT) and various mutant strains was digested with specific restriction enzymes and hybridized with probes targeted to internal or external loci. For *tea1-1*, *tea1-2*, *tea5-2*, *chs1-1*, *chs1-2*, *chs1-4*, *sep4*, and *sep5*, the blots exhibit clear and distinct bands corresponding to the predicted sizes for the replacement of the open reading frame with the selection marker. For *sep3*, *sep6*, *nox1-1-*, *nox1-2*, and *noxr1*, the blots display multiple bands, suggesting the occurrence of ectopic insertions elsewhere in the genome; however, the complete absence of the specific WT band in these strains confirms that the endogenous target genes were successfully disrupted.

**Figure EV6.**
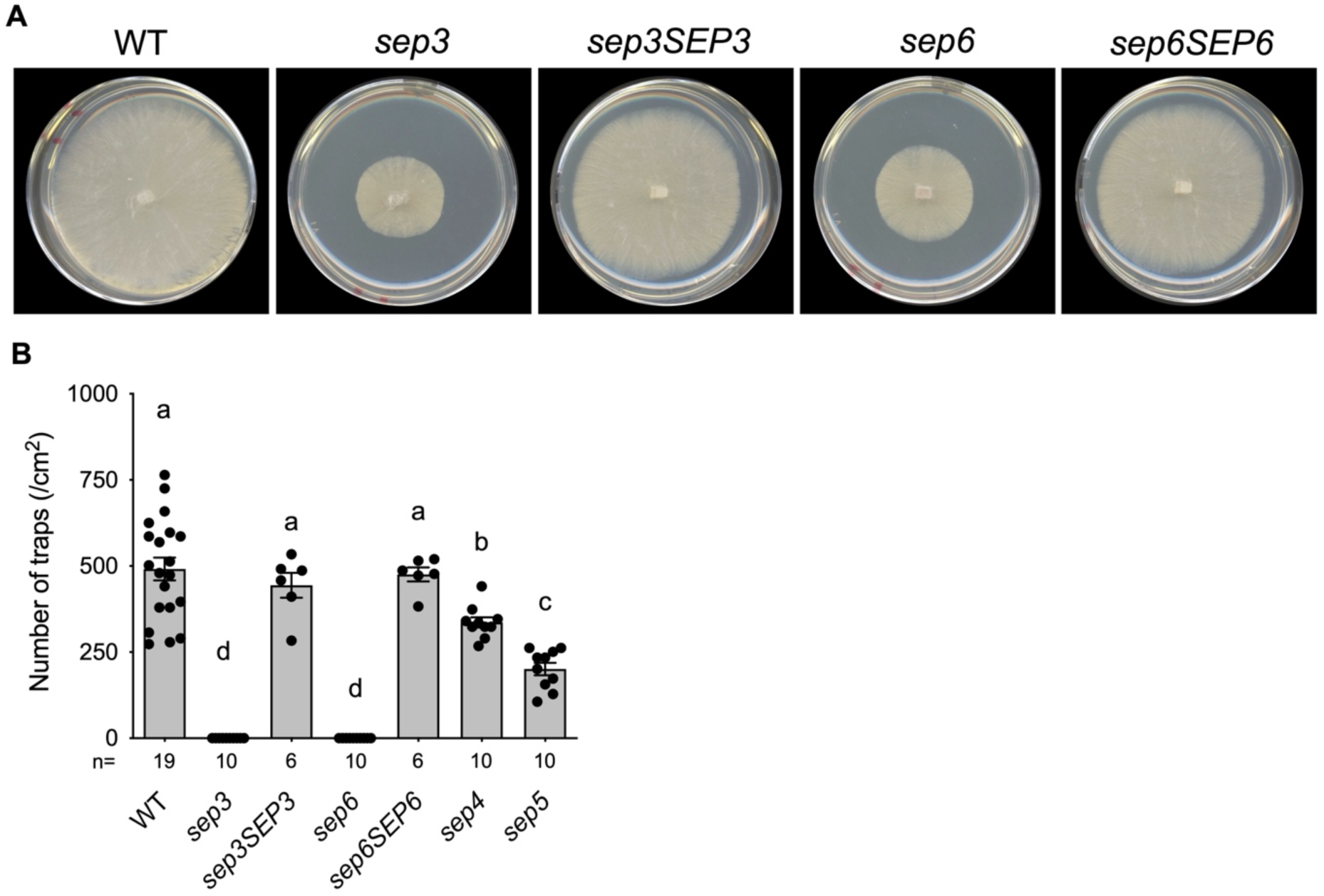
Sep3 and Sep6 are required for fungal growth and trap formation. **(A)** Colonies of the wild type (WT), *sep3*, *sep3SEP3*, *sep6* and *sep6SEP6* strains grown on 5.5-cm PDA plates for 3 days. **(B)** Quantification of the trap numbers induced by 30 *C. elegans* larvae for the WT, and *sep* mutant lines. Data represent mean ± SEM; n is shown along the x axis; different letters indicate significant differences based on one-way ANOVA followed by uncorrected Fisher’s LSD multiple test, with a threshold of *P* < 0.05.

